# Self-reported sleep problems are related to cortical thinning in aging but not memory decline and amyloid-β accumulation – results from the Lifebrain consortium

**DOI:** 10.1101/2020.04.28.065474

**Authors:** Anders M. Fjell, Øystein Sørensen, Inge K. Amlien, David Bartrés-Faz, Andreas M. Brandmaier, Nikolaus Buchmann, Ilja Demuth, Christian A Drevon, Sandra Düzel, Klaus P. Ebmeier, Paolo Ghisletta, Ane-Victoria Idland, Tim C. Kietzmann, Rogier A. Kievit, Simone Kühn, Ulman Lindenberger, Fredrik Magnussen, Didac Macià, Athanasia M. Mowinckel, Lars Nyberg, Claire E. Sexton, Cristina Solé-Padullés, Sara Pudas, James M. Roe, Donatas Sederevicius, Sana Suri, Didac Vidal-Piñeiro, Gerd Wagner, Leiv Otto Watne, René Westerhausen, Enikő Zsoldos, Kristine B. Walhovd, for the Alzheimer’s Disease Neuroimaging Initiative

**Author notes:** Address correspondence to: Anders M Fjell, Dept of Psychology, Pb. 1094 Blindern, 0317 Oslo, Norway. Some of the data used in preparation of this article were obtained from the Alzheimer’s Disease Neuroimaging Initiative (ADNI) database (adni.loni.usc.edu). As such, the investigators within the ADNI contributed to the design and implementation of ADNI and/or provided data but did not participate in analysis or writing of this report. A complete listing of ADNI investigators can be found at: http://adni.loni.usc.edu/wp-content/uploads/how_to_apply/ADNI_Acknowledgement_List.pdf as well as in the Supplementary material.

## Abstract

**Background:** Older persons with poor sleep are more likely to develop neurodegenerative disease, but the causality underlying this association is unclear. To move towards explanation, we examine whether sleep quality and quantity are similarly associated with brain changes across the adult lifespan.

**Methods:** Associations between self-reported sleep parameters (Pittsburgh Sleep Quality Index;PSQI) and longitudinal cortical change were tested using five samples from the *Lifebrain* consortium (n=2205, 4363 MRIs, 18-92 years). Analyses were augmented by considering episodic memory change, gene expression from the Allen Human Brain Atlas, and amyloid-beta (Aβ) accumulation (n=1980).

**Results:** PSQI components sleep problems and low sleep quality were related to thinning of the right lateral temporal cortex. The association with sleep problems emerged after 60 years, especially in regions with high expression of genes related to oligodendrocytes and S1 pyramidal neurons. BMI and symptoms of depression had negligible effects. Sleep problems were neither related to longitudinal change in episodic memory function nor to Aβ accumulation, suggesting that sleep-related cortical changes were independent of AD neuropathology and cognitive decline.

**Conclusion:** Worse self-reported sleep in later adulthood was associated with more cortical thinning in regions of high expression of genes related to oligodendrocytes and S1 pyramidal neurons, but not to Aβ accumulation or memory decline. The relationship to cortical brain change suggests that self-reported sleep parameters are relevant in lifespan studies, but small effect sizes, except for a few restricted regions, indicate that self-reported sleep is not a good biomarker of general cortical degeneration in healthy older adults.

## Introduction

Older adults with poor sleep are more likely to develop neurodegenerative disease^1-8^, and in a recent meta-analysis it was argued that 15% of Alzheimer’s Disease (AD) diagnoses in the population may be attributed to aspects of sleep^9^. It is not known whether this relationship points to a causal connection between sleep and neurodegeneration, and little is known about the mechanisms that could mediate such a connection. For instance, the association might be bidirectional or reflect the operation of mechanisms (e.g.^10^) affecting both sleep problems and brain change in normal and pathological aging. Change in sleep patterns is a feature of normal aging^11-13^, so it is important to know whether natural variations in sleep quality and quantity are related to brain changes across adult life. Here, we examined the relationship between multiple self-reported sleep variables, including sleep problems and sleep duration, and cortical changes across the adult lifespan. Participants from the Lifebrain consortium assessed repeatedly (up to 7 assessments over 11 years) with MRIs were studied^14^. In additional analyses we tested coherence with cell-specific gene expression maps from the Allen Human Brain Atlas, and with changes in memory performance and amyloid-beta accumulation (Aβ).

Several studies have reported cross-sectional relationships between self-reported sleep parameters and structural properties of the cerebral cortex. Poor sleep tends to be associated with thinner cortex or smaller cortical volume (Table 1), but with little agreement about the self-reported sleep measures and cortical regions involved. The largest study to date found daytime sleepiness to be associated with thinner cortices in all lobes^15^, but did not report results for other sleep variables. Three studies tested associations with longitudinal cortical change. In one, lower global sleep quality was related to greater cortical volume reduction widespread in frontal, temporal, and parietal regions^16^ in participants above 60 years of age. Sleep duration was not associated with atrophy, in accordance with a recent study where duration measured five times over 28 years^17^ in 613 participants did not relate to global or regional gray matter volume measured at follow up. In contrast, a second study found that sleep duration of more or fewer than 7 hours was associated with frontotemporal cortical thinning^18^. A third study of middle-aged and older adults reported shorter sleep duration to be related to more ventricular expansion, whereas no relationship was observed between a global sleep score and volumetric change in total cerebral, inferior or superior frontal volume^19^. Thus, poor self-reported sleep tends to be modestly related to unfavorable cortical changes, but with little consistency across studies with regard to sleep parameters and brain regions.

*[Table 1]*

Here we took advantage of five longitudinal MRI studies from the Lifebrain consortium^14^ to examine which self-reported sleep parameters were related to cortical changes and whether these associations were stable through adult life or increased in strength with age. We hypothesized stronger relationships between sleep and cortical changes in older compared to younger adults, mainly in frontal and temporal cortices. These hypotheses were speculative due to few longitudinal studies and highly divergent findings in previous publications. In additional analyses we tested whether sleep and cortical thinning were preferentially associated in regions characterized by high rates of natural age changes, whether such relationships were stronger in cortical regions characterized by higher expression of genes related to specific cell types – so-called “virtual histology”^20^. Guided by results from animal experiments, overlap with regions of high expression of microglia, astrocytes and oligodendrocyte genes could be expected. Given that a link between poor sleep and the AD biomarker Aβ has been proposed^21-25^, we also tested the association between sleep problems and Aβ in samples of cognitively normal older adults (total n = 264), as well as in 1716 Alzheimer’s Disease Neuroimaging Initiative (ADNI; adni.loni.usc.edu) participants across the AD spectrum. We hypothesized a positive relationship between poor sleep and Aβ. We also examined whether there was a relationship with memory decline

## Methods and materials

### Sample

The sample was derived from the European Lifebrain project

(http://www.lifebrain.uio.no/)^14^, including participants from the Berlin Aging Study II (BASE-II)^26,27^, the BETULA project^28^, the Cambridge Centre for Ageing and Neuroscience study (Cam-CAN)^29^, the Center for Lifespan Changes in Brain and Cognition longitudinal studies (LCBC)^30,31^, and the University of Barcelona brain studies^32-34^. Self-reported sleep and structural MRIs from 2205 participants (18.5-91.9 years) yielding a total of 4363 observations were included (Table 2). Screening criteria were not identical across studies, but participants were in general cognitively healthy and did not suffer from neurological conditions known to affect brain function, such as dementia, major stroke and multiple sclerosis (Fjell et al.^35^ and SI). Depressive symptoms were mapped using the Beck Depression Inventory^36^ (LCBC, except MADRS for n=91), the Geriatric Depression Scale^37^ 15 item version (BASE-II), the Center for Epidemiologic Studies Depression Scale^38^ (Betula), and the Hospital Anxiety and Depression Scale^39^ (Cam-CAN).

Relationship between sleep parameters and Aβ were tested in samples of cognitively normal older adults (n = 264) (Table 3) (see SI). Aβ accumulation was measured by CSF Aβ42, ^18^F-flutemetamol or ^18^F-florbetapir PET. Separate analyses were conducted in two additional samples from ADNI, where sleep problems were measured by NPI-Q^40^, of normal controls (NC, n = 655; 1423 observations), patients with mild cognitive impairment (MCI, n = 765; 1541 observations) and AD (n = 296; 690 observations). Aβ was quantified from ^18^F-florbetapir PET (n = 846) or CSF Aβ42 (n = 870). As most ADNI participants had multiple Aβ measurements, participants with more CSF than PET observations were placed into the ADNI CSF dataset and vice-versa. Participants with equal number of CSF and PET observations were randomly assigned to either the CSF or PET dataset.

Associations between sleep parameters and longitudinal memory change were tested in 1419 participants (2702 observations; Betula [n = 138/ 276 observations, baseline-age = 65, 55-85 years], LCBC [n = 730/ 1480 observations, baseline-age = 49, 19-89 years], BASE-II [n = 512/ 830 observations, baseline-age = 62, 24-92 years], Barcelona [n = 39/ 116 observations, baseline-age = 69, 64-81 years]), by 30 min delayed free recall from the Verbal Learning and Memory Test (BASE-II), 30 min free recall from the California Verbal Learning Test (LCBC)^41^, word recall (immediate free recall of words and sentences, and delayed cued recall of words from the previously learned sentences, Betula)^42,43^ and 30 min delayed recall from the Rey Auditory Verbal Learning Test ^44^ (Barcelona).

### Self-reported sleep assessment

Sleep was assessed using the Pittsburgh Sleep Quality Inventory (PSQI)^45^ representing self-reported sleep quality for the previous month. This assessed 7 domains (sleep quality, latency, duration, efficiency, problems, medication and daytime tiredness) each scored from 0 to 3 and a global score. High scores indicate worse sleep. Duration was also tested using number of hours instead of the 0-3 scale. The sleep medication item was not analyzed, as most samples used screening for use of medications. The Karolinska Sleep Questionnaire (KSQ)^46,47^ was used for Betula and converted to PSQI (SI and ^35^). Although lifespan changes in sleep are pronounced^35,48^, mean changes as well as individual differences in change over a few years are rather small^17,49^ and considered as negligible here. If multiple observations were available, the mean value was used.

### Magnetic resonance imaging (MRI) acquisition and analysis

MRI data originated from 8 different scanners (Table 4), processed with FreeSurfer v6.0 (https://surfer.nmr.mgh.harvard.edu/)^50-53^, yielding maps of cortical thickness and volume. To avoid introducing possible site-specific biases, quality control measures were imposed and no manual editing was done. We previously have reported that across-scanner consistencies in estimated hippocampal volumes for the scanners used in Lifebrain are high. We nevertheless included ‘study site’ as a fixed effect covariate to adjust for scanner differences^35^.

### Virtual histology analyses – relationship with cell-specific gene-expression profiles

We tested how sleep-associated cortical thinning related to regional gene expression profiles associated with specific cell types. As described in detail elsewhere^20,54^, gene-expression data in *ex-vivo* left hemisphere brains were obtained from the Allen Human Brain Atlas (AHBA: (http://www.brain-map.org^55^), mapped to 34 cortical regions of the Desikan-Killiany Atlas^56^. Genes were required to be regionally consistent across 6 donors and two datasets (AHBA, the BrainSpan Atlas) (SI and^54^). The resulting 2511 genes with consistent expression profiles were combined with a list of cell-specific genes from Zeisel et al.^57^, yielding the following cell-type panels: S1 pyramidal neurons (n=73), CA1 pyramidal neurons (n=103), interneurons (n=100), astrocytes (n=54), microglia (n=48), oligodendrocytes (n=60), ependymal (n=84), endothelial (n=57), and mural (n=25).

### Statistical analyses

See SI for details.

#### Cortical surface analyses

Spatiotemporal linear mixed models^58,59^ implemented in FreeSurfer v6.0.1 and running on MATLAB R2017a were used to test relationships between sleep variables and cortical thickness/ volume change, accounting for the spatial correlation between residuals at neighboring vertices and the temporal correlation of residuals within repeated measurements of single participants. Surface results were tested against an empirical null distribution of maximum cluster size across 10 000 iterations using Z Monte Carlo simulations, yielding results corrected for multiple comparisons across space (p< .05) ^60^. Baseline age, time since baseline, site and sex were included as covariates. Separate analyses included all two- and three-way interactions between PSQI, age, and time. The hierarchical nature of the data (repeated measurements nested within participants) was accounted for using a random intercept term. Additional analyses were conducted, using a dichotomous indicator variable for age (≥ vs. < 60 years), creating a piecewise linear model for cortical thinning with a cut-point at 60 years. All follow-up analyses were run in R^61^. To assess relationships between PSQI and memory change, we ran generalized additive mixed models^62^ (GAMM) with a random intercept and random effects per study for time and sleep using the package “mgcv” using smooth terms for baseline age. Additional models were run with PSQI×time×baseline-age and age×sleep.

#### Virtual histology

Pearson correlations were computed between the thinning coefficients and the median inter-regional profile of gene expression levels for each marker gene. Using a resampling-based approach (Shin et al., 2018b), the average expression - thinning correlation for each group of cell-specific genes served as the test statistic. False discovery rate was used to account for 9 cell-types.

#### Aβ-sleep

The association between sleep and Aβ was tested by partial correlations, controlling for age and sex. We performed additional analyses in ADNI by generalized linear mixed effect regressions (logistic link function with binominal error distribution; “lme4” R-package). Sleep disturbance (dichotomous) was outcome, clinical diagnosis, Aβ, age and sex were fixed effects and subjects were random effects. Statistical significance was assessed with Kenward-Roger-corrected tests as implemented with the “car” R-package^63^.

## Results

### Worse self-reported sleep is associated with accelerated cortical thinning

Associations of PSQI scales and cortical atrophy are shown in Figure 1 and Table 5. Lower sleep quality was related to more cortical thinning in the right middle temporal gyrus, extending posteriorly. A similar relationship for sleep problems was found in an overlapping region, bordering angular gyrus. For sleep problems, extensive age-interactions were found (Figure 2), with three large lateral clusters in the right hemisphere, located in the temporal, inferior parietal and inferior/ middle prefrontal cortex, and one cluster in the left (lateral, inferior and medial temporal cortex). More sleep problems were related to accelerated cortical thinning from about 60 years of age (see below). For the remaining PSQI components and the global score, no effects were found. No relationships with cortical volume change were seen. Follow-up analyses were performed to test whether sleep duration showed a quadratic relationship with atrophy. Here, duration was measured in hours instead of the PSQI 0-3-point scale. There were no significant quadratic relationships or age-interactions.

**Figure 1.**
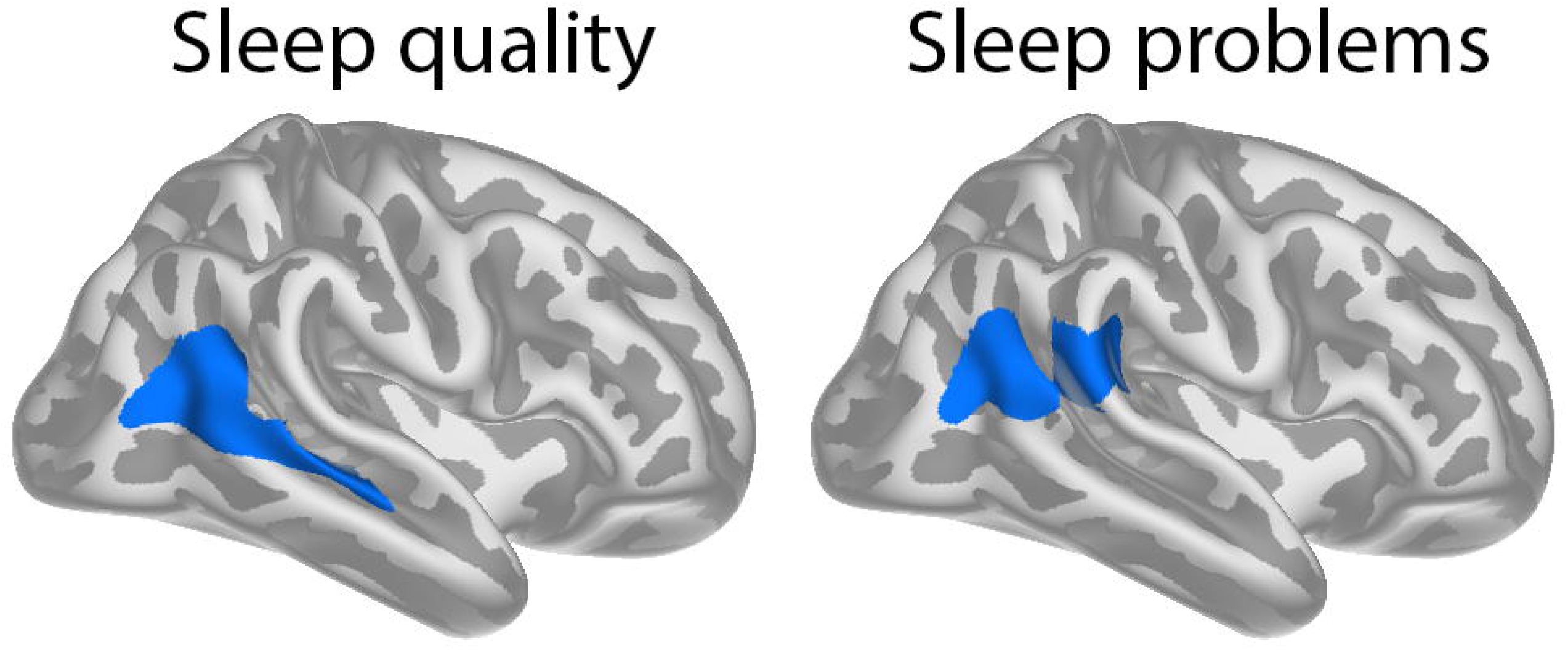
Effects of sleep on cortical thinning. Clusters where lower self-reported sleep quality and more sleep problems are related to cortical thinning after corrections for multiple comparisons across space.

**Figure 2.**
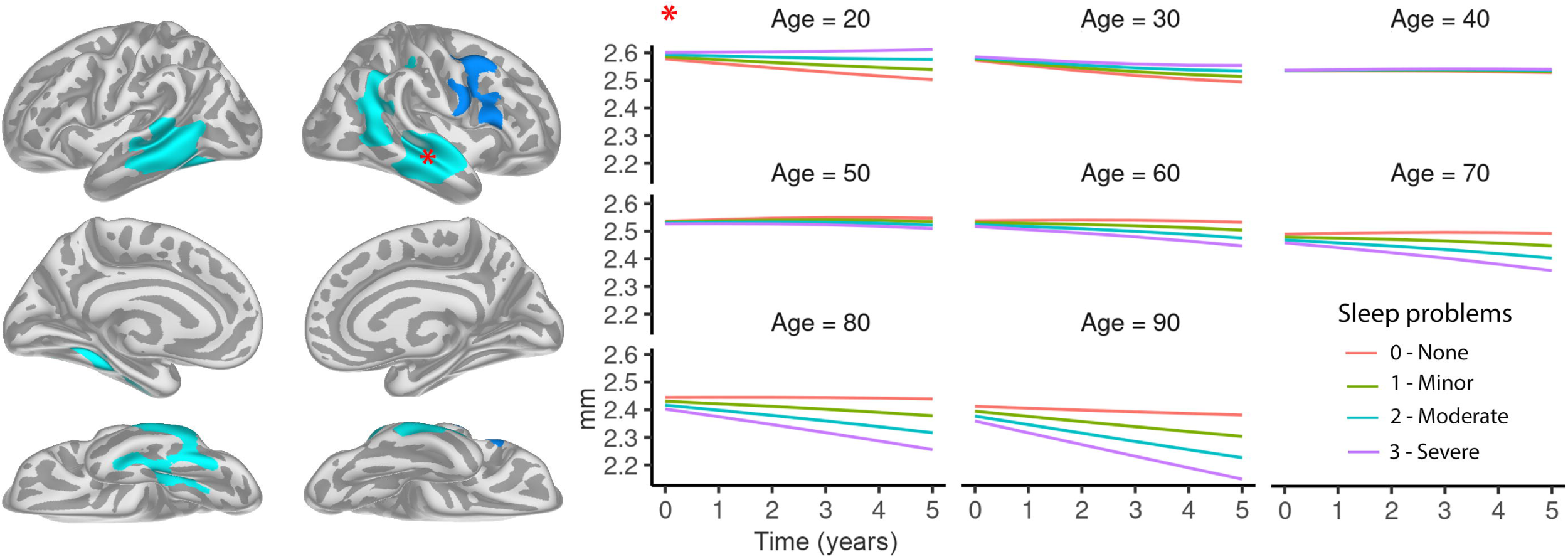
Age interactions Left panels: Clusters where more self-reported sleep problems are significantly more strongly related to cortical thinning in older than younger adults. Right panels: Model predicted cortical thickness in the right middle temporal cluster in the left panel as a function of age, time and amount of sleep problems (0: none, 3: severe). Participants reporting more sleep problems show more cortical thinning in the older age-ranges but not in the younger. CIs round the curves were removed for improved viewing, and are presented in SI, along with similar curves for the other three significant clusters. These plots are used for illustrating the effects in the surface analyses. Statistical analyses were conducted using age as a continuous variable.

As can be seen in Figure 2, the association between sleep problems and cortical thinning emerged from around 60 years of age. To assess the size of this age interaction, we calculated mean cortical thinning (Δthickness mm/year) as a function of sleep problems in 60-year olds and 80-year-olds (Figure 3). For participants with minor sleep problems, annual cortical thinning was low and close to identical for 60-year olds and 80-year olds; ≈0.05 mm/year or -0.20%. In contrast, for participants with severe sleep problems, annual cortical thinning was almost twice as large in the oldest group, equal to -1.03% (0.027 mm/year) compared to -0.55% (0.014 mm/year), respectively.

**Figure 3.**
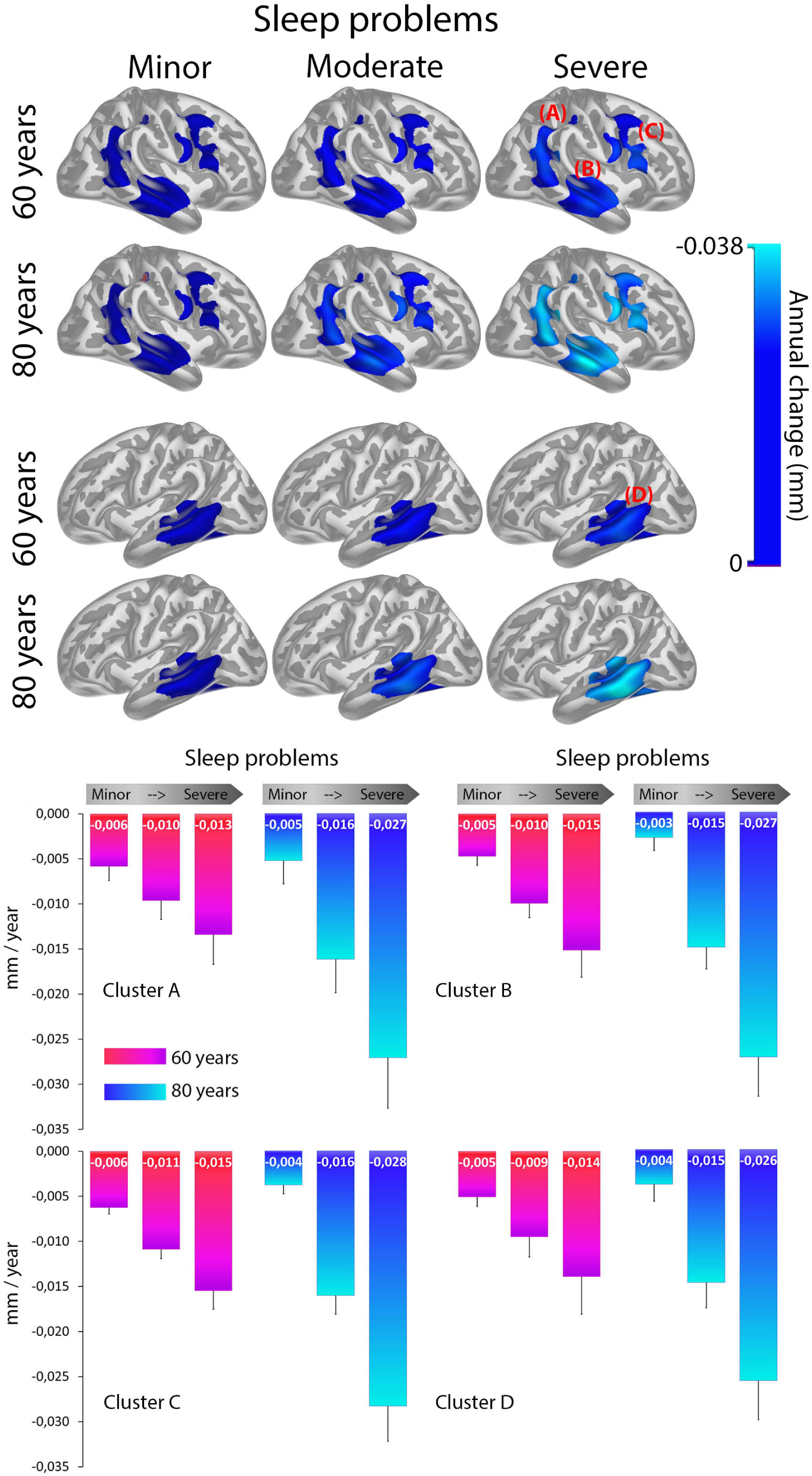
Cortical thinning as a function of sleep problems and age Top panel: Mean cortical thinning (mm/year) in the clusters with a significant effect of age × sleep on thickness change broken up by degree of sleep problem and age group. Bottom panel: Mean cortical thinning in the same regions as a function of sleep problems and age group. Error bars denote 1 SD.

### Sleep problem-related cortical thinning is not stronger in regions showing the most atrophy

To test whether the age-interactions for sleep problems were strongest in regions showing the most atrophy in aging, we calculated the vertex-wise spatial correlations between the age × sleep problems interaction coefficients and cortical thinning in participants above 60 years. The correlations were .04 for the left and -.003 for the right hemisphere. These very low correlations suggest that sleep problems are not more related to cortical thinning with higher age in the regions undergoing the most thinning per se, although bootstrapping showed that the left hemisphere relationship was significantly larger than expected by chance (p < .001).

### Sleep-related thinning and expression of oligodendrocytes and S1 pyramidal cell genes

Virtual histology was used to test whether the age-dependent associations between sleep problems and cortical thinning were related to inter-regional gene expression profiles associated with specific cell types (Table 6, Figure 4). The results showed overlap with regions with higher expression of genes related to oligodendrocytes and S1 pyramidal cells (P_fdr_ < .05). Note that these analyses were performed vertex-wise, and were not restricted to regions with significant sleep×time×age interactions.

**Figure 4.**
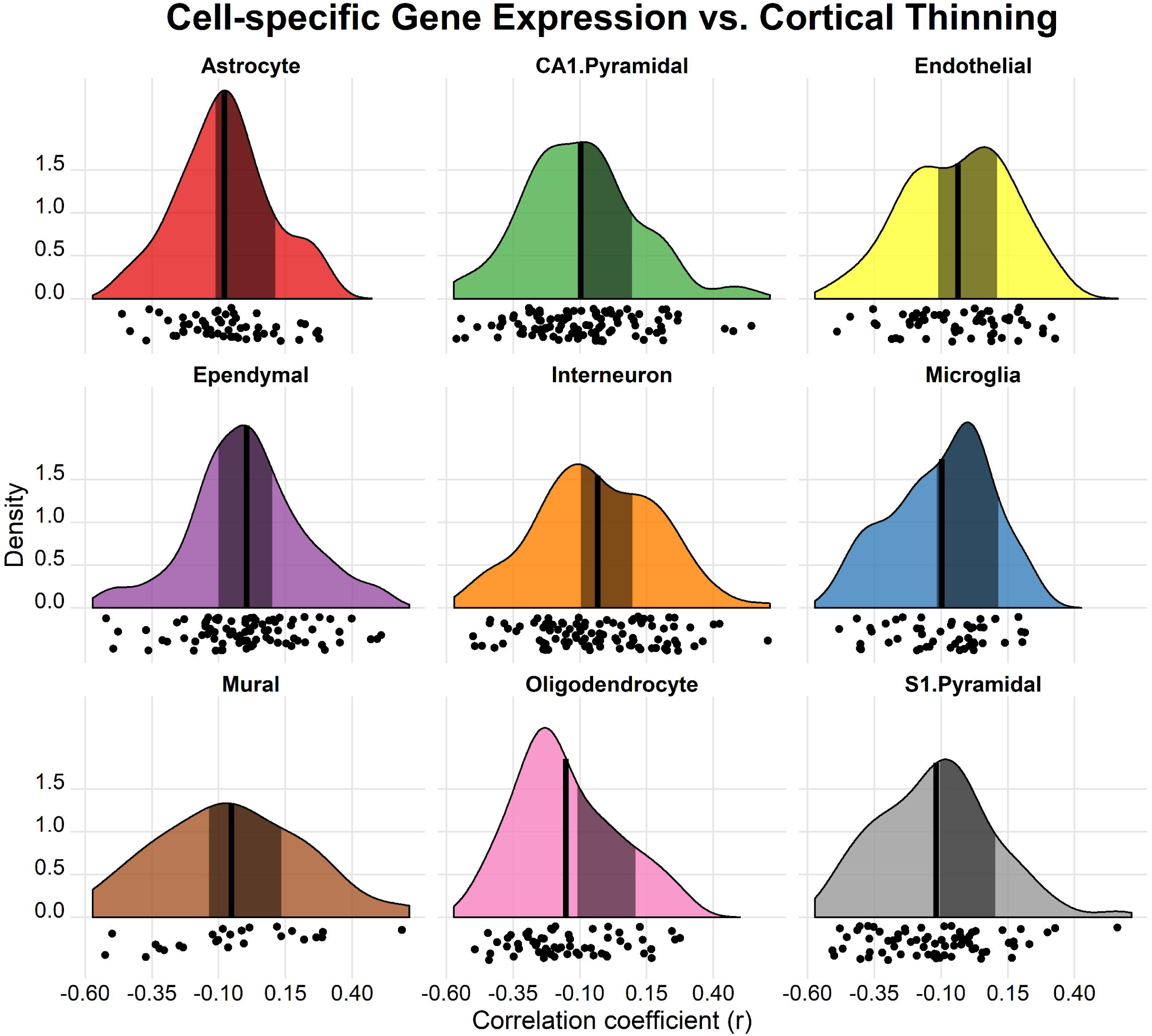
Virtual histology. Significantly higher expression of genes characteristic of specific cell types (oligodendrocytes and S1 pyramidal) in vertices where the age-related association between sleep problems and cortical thinning are the strongest (vertex-wise thickness effects from the age × time × sleep interaction; see surface plots in Figure 2). The black vertical lines represent the mean association, and the shaded area represents the empirical null distribution for each of the 9 cell types. The x-axes indicate the coefficients of correlation between the thinning and the gene expression profiles. The y-axes indicate the estimated probability density for the correlation coefficients. Panels where the mean of the target ROIs is outside the null distribution 95% C.I. are considered to show a correlation greater or smaller than predicted. See ^92^ for creation of raincloud plots.

### No relationship with Aβ accumulation

First, we tested the correlations between sleep problems and biomarkers of Aβ accumulation in each of the Lifebrain subsamples, controlling for age and sex. Correlations between Aβ and sleep problems were r = .11 (p = .15, n = 173) and r = .03 (p = .80, n = 91) for the PET and CSF subsamples, respectively. We performed additional analyses in the ADNI cohort of NC, MCI and AD, using linear mixed effects logistic regression analyses. Neither for the PET (p = .86) or the CSF ADNI sample (p = .46) was Aβ related to sleep problems, whereas there was a significant effect of diagnosis with more sleep problems in patients compared to normal controls (PET sample Chi sq = 25.59, p < 1.7e^-6^; CSF sample Chi sq = 18.38, p < .00028). Although done on a different type of sample, the ADNI results concur with the results from the Lifebrain Aβ-samples.

### No relationship with memory change

GAMMs were performed with memory score as outcome, PSQI × time as the main term of interest, with sex and baseline age as covariates. No significant relationships were found (all p’s ≥ .22). Effect estimates are presented in SI, showing that the largest effect was found for sleep efficiency, for which the efficiency (scale 0-3) × time (in years) interaction on change in memory scores (SD) was -0.022, i.e. the lowest sleep efficiency (“0”) was associated with 0.33 SD relative decline in memory score over a five-years interval compared to the highest efficiency (“3”). We tested the three-way interaction PSQI×time×baseline-age, with no significant effects (all p’s ≥ .58). Finally, we ran models with the smooth interaction between age at testing and PSQI. Here the longitudinal and cross-sectional information is combined in the test age variable, and because chronological age has much more variance than ages at repeated measures, this analysis is skewed towards cross-sectional effects. Here, age×sleep problems showed a weak but significant relationship with memory change (p = .039, uncorrected). None of the other PSQI components showed this, with duration (p = .086) and efficiency (p = .099) scales yielding the lowest p-values, and global score the highest (p = .96).

### Post hoc analyses: controlling for BMI and depression

We performed post hoc analyses testing whether body mass index (BMI) or preclinical depression affected the observed sleep problem – cortical change relationship (see SI for details). BMI was available at the time of analysis from the LCBC, CamCAN and Betula datasets (n = 1717), and depression scores from the LCBC, CamCAN, Betula and BASE-II datasets (n = 1637). As seen in Figure 5 and 6, controlling for BMI or depression symptoms did not affect the relationship between sleep problems and cortical change, except for some offset effects for BMI in the 50ties and 60ties, not affecting the actual slope.

**Figure 5.**
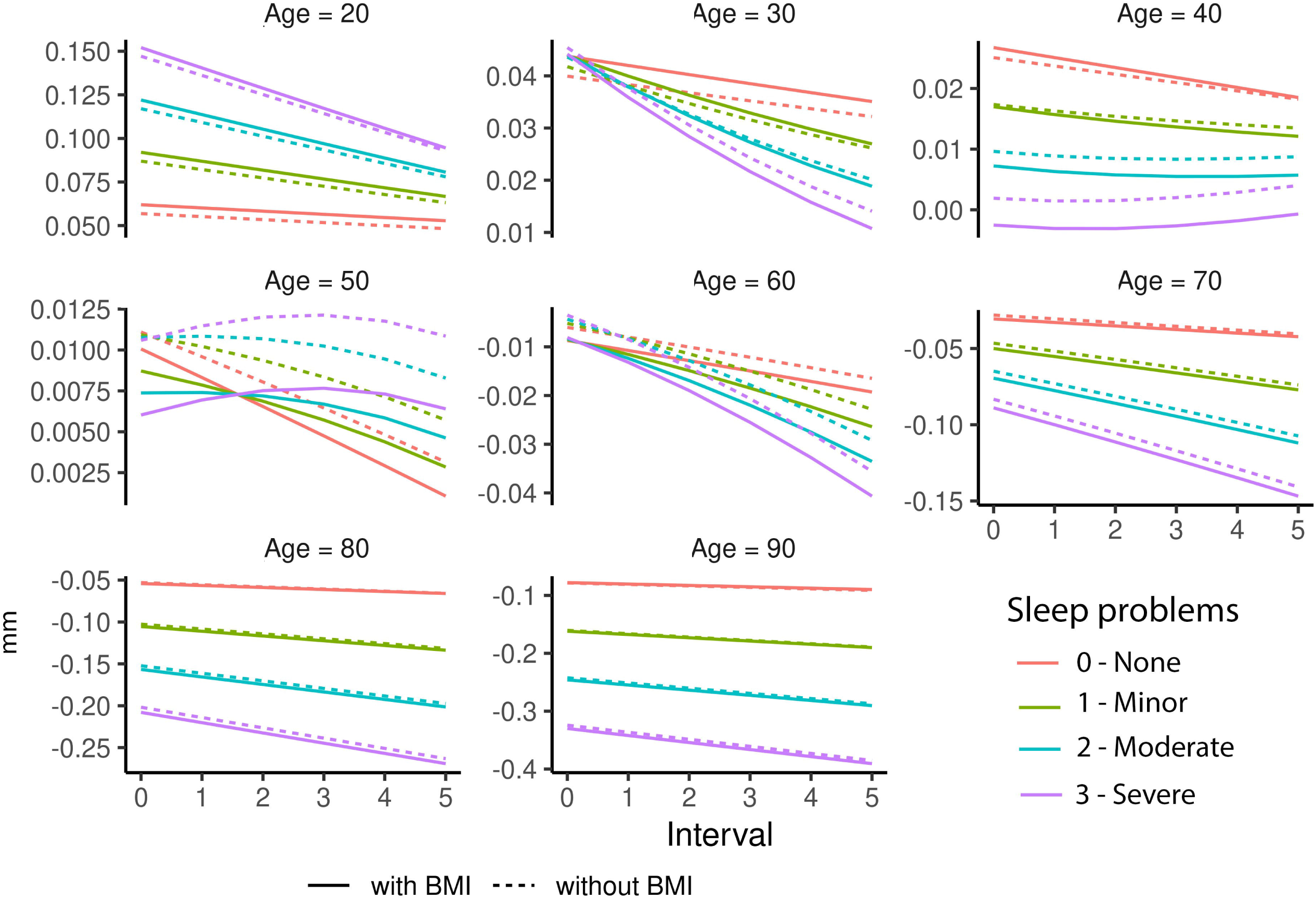
Controlling for BMI. BMI did not affect the relationship between sleep problems and cortical thinning across age, as can be seen from the overlapping solid and dotted lines. Plotted effects are from Cluster B (Figure 3); see SI for additional clusters. Please note that the y-axes are fitted to each plot to facilitate detection of possible differences between curves visually, and therefore vary between plots.

**Figure 6.**
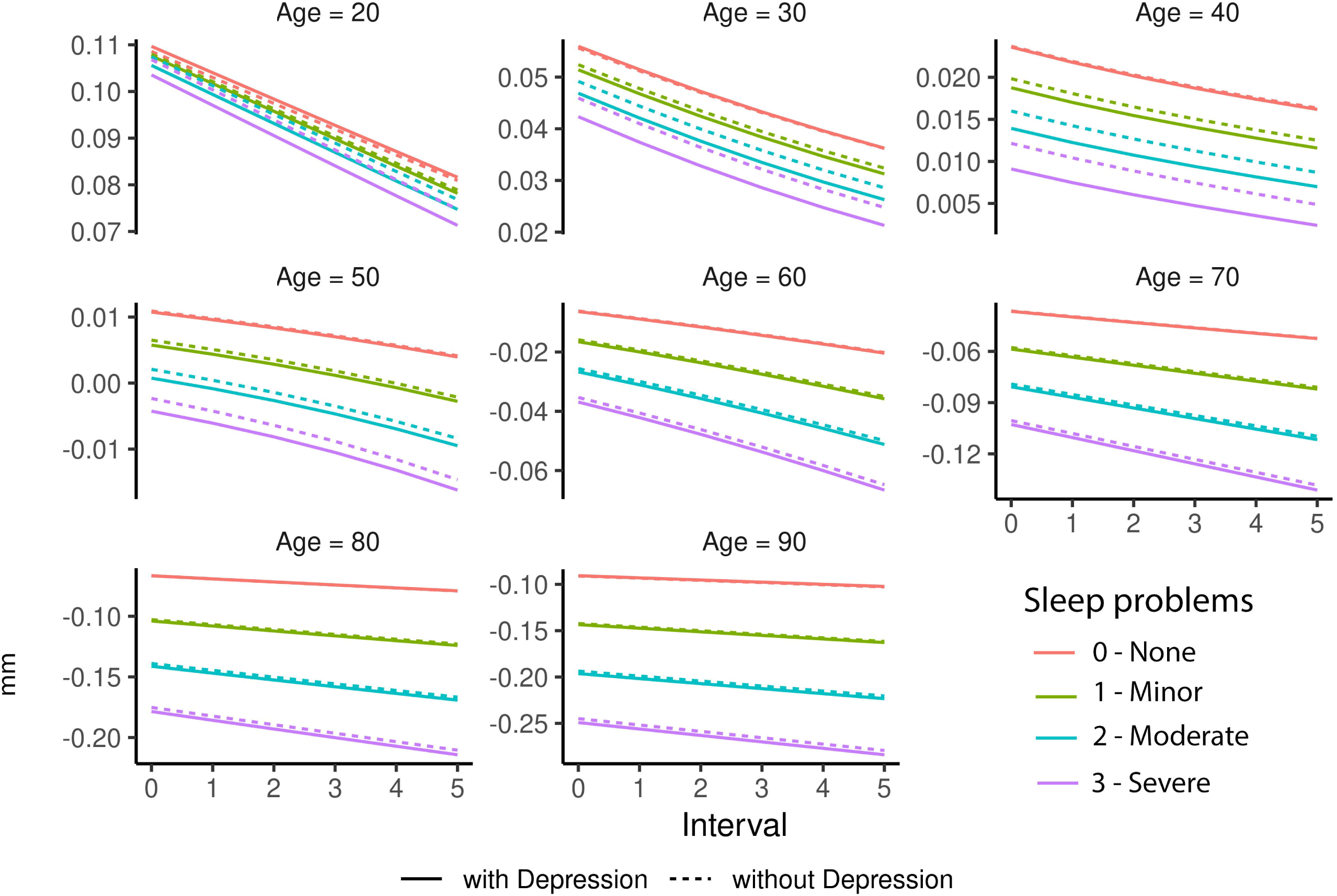
Controlling for depression. Depressive symptoms did not affect the relationship between sleep problems and cortical thinning across age, as can be seen from the overlapping solid and dotted lines. Plotted effects are from Cluster B (Figure 3); see SI for additional clusters. Please note that the y-axes are fitted to each plot to facilitate detection of possible differences between curves visually, and therefore vary between plots.

## Discussion

Lower self-reported sleep quality and more sleep problems were associated with more cortical thinning, independently of depressive symptoms and BMI, after the age of 60 years. These relationships were stronger in regions with higher expression of genes related to oligodendrocytes and S1 pyramidal cells. The lack of relationship with Aβ accumulation suggests that this is not dependent on preclinical AD pathology. At the same time, effect sizes were modest, there were no significant associations with global sleep quality or sleep duration, and there was no significant association with memory change.

### Sleep problems and cortical thinning

Lower sleep quality and more sleep problems were related to more cortical thinning in the right middle temporal gyrus, extending posteriorly. Sleep quality is based on the participants’ overall evaluation of their sleep, whereas the sleep problem scale includes items such as nocturnal awakenings, bad dreams and trouble breathing. In previous work on a subsample of the current sample, we found global sleep problems to be related to volume reductions in overlapping regions^16^ in older adults. Although sleep problems may accelerate cortical atrophy in aging, stronger relationships were not found in the regions that showed most cortical thinning. This suggests that sleep problems do not simply relate to general patterns of cortical atrophy that are typical in aging, but more regionally specific patterns of cortical decline. Alternatively, these results, together with previously reported unspecific relationships between sleep, cognition and neurodegeneration^11-13^, suggest that sleep problems result from multiple changes in the aging brain. Sleep and brain decline are probably linked through complex interactions with multiple other factors, some involving atrophy and some not^31^.

It is important to stress that effect sizes were small. Even with 4363 MRIs, only two aspects of self-reported sleep were significantly associated with change in cortical thickness. Thus, the standard self-report sleep metrics do not seem to have much clinical relevance as markers of cortical atrophy. For participants in their sixties, the transition from minor to severe sleep problems was weakly but reliably associated with cortical thinning. For 80-years old participants, that same transition was accompanied by a rather substantial increase in cortical thinning. However, as participants were not followed over time with repeated sleep measures, we do not know whether these group effects translate to relationship within individuals. Thus, although our findings are interesting as they demonstrate an age-dependent relationship between sleep problems and cortical thinning, the mechanisms driving this age dependency in the strength of the association between sleep problems and cortical thinning await further study.

### Virtual histology – a possible role of oligodendrocytes

The age-dependent sleep association overlapped with regions characterized by higher expression of genes related to oligodendrocytes and S1 pyramidal cells. These results do not reveal any direct mechanistic relationships but yield useful information for interpretation of the findings, especially regarding genes related to oligodendrocytes. Several genes involved in the synthesis and maintenance of myelin are expressed at higher levels during sleep, and sleep loss may negatively impact oligodendrocyte physiology^64^. Specifically, genes involved in phospholipid synthesis and myelination tend to be transcribed preferentially during sleep^65^. For instance, gene expression analysis performed on oligodendrocyte-enriched samples of mouse forebrain showed that several genes changing their expression in sleep and awake were oligodendrocyte precursor cell specific genes^65^. Genes promoting oligodendrocyte precursor cell (OPC) proliferation were mainly upregulated during sleep whereas OPC differentiation were mainly upregulated during awake. Such studies suggest that sleep is important for myelination^66^. Myelin properties around the GM/WM boundary affect cortical thickness estimates^67,68^ and change through the lifespan^69,70^. Thus, regions with higher expression of oligodendrocyte-related genes could be more vulnerable to sleep problems, explaining the overlap between expression of oligodendrocyte-related genes and sleep-age-related cortical thinning. Although speculative, experimental and large-scale genetic studies could address this.

### No relationship between cortical thinning, sleep duration, memory change and Aβ

Sleep duration and cortical thinning was not associated. Sleep duration is the most frequently investigated sleep measure in relation to health^71^, and is possibly modifiable. It has been suggested that we have a ‘global epidemic of sleeplessness’^72,73^, and the United States National Sleep Foundation (NSF) provides sleep duration recommendations^74,75^. Previous studies have not provided conclusive evidence about a role for sleep duration in brain change^16,18,19,35^. We found no indication that sleep duration was related to cortical change, supporting previous Lifebrain results on hippocampal atrophy^35^. In that study, however, participants from the UK Biobank reporting short (< 5 hours) and long (> 9 hours) sleep duration showed smaller hippocampal volume. Although we cannot conclude with confidence that extremely short or extremely long sleep duration is unrelated to cortical atrophy, our results demonstrate that within normal ranges of sleep duration, there is no association with cortical change.

Sleep problems were not related to changes in memory function or Aβ accumulation. Although a relationship between Aβ and sleep has been suggested from observational^21-24^ and experimental human studies^25^, the association is not clear across reports. One study found sleep latency but not problems to be related to Aβ^21^, another found relationships to latency and quality, but not number of nocturnal awakenings, which would be most similar to the measure of sleep problems^22^. A third study found sleep problems together with two other self-report measures to be related to more Aβ, whereas four other measures were not^23^, a fourth reported a relationship for less adequate sleep but not problems and two additional measures^76^, and a fifth a relationship with global sleep quality in participants below but not above 67 years^77^. Thus, a robust relationship between sleep problems and Aβ accumulation has not been established. Here, with close to 2000 participants, we found no relationship with sleep problems.

Similarly, we did not observe a relationship between sleep problems and memory change. The only significant result was the interaction between sleep problems and age, which would not survive correction for multiple comparisons. Experimental studies often report negative effects of short-term sleep restriction on cognition^78^, but it is unclear whether such effects are relevant in the long run and in a naturalistic setting. Meta-analyses have reported poorer episodic memory in people with insomnia^79^, whereas epidemiological studies have yielded mixed results and the largest study to date – based on 477.529 participants from the UK Biobank – found no relationship between insomnia and any cognitive function^80^. The possibility remains that longitudinal changes in self-reported sleep could be related to declining memory over time^31^, and stronger relationships could potentially be seen with other measures of sleep^81,82^. Still the present results, based on 1419 participants and 2702 memory tests, suggest that if a relationship between self-reported sleep and memory decline does exist, the effect size of this association can be expected to be small.

### Limitations

Causes and appearance of age-related changes in sleep are manifold and diverse^83^. There is no perfect way to measure sleep without disrupting routine^84^. Self-reports of sleep duration are only moderately correlated with actigraph measures^84-86^, whereas actigraphs themselves tend to over-estimate sleep duration compared to polysomnography^87-91^. Still, self-report measures continue to be important in sleep studies, and epidemiological studies and the NSF recommendations are primarily based on self-reports^74^. Although we acknowledge the uncertainty of self-reports, this is currently the only viable option for testing sleep-brain relationships in large samples. Also, we are mot able to distinguish between effects of short- vs. long-term sleep patterns, as PSQI only index sleep over the last month. Another possibe limitation is that data were pooled across studies using partly different measures. Whereas this may introduce noise, it also reduces the risk of non-representative and biased samples and greatly increases statistical power.

## Conclusion

Worse self-reported sleep was related to increased rate of cortical thinning after the age of 60 years in regions with higher expression of genes related to oligodendrocytes and S1 pyramidal cells. This makes self-reported sleep a relevant variable in studies of brain aging, independently of AD pathology. Small effect sizes and lack of relationship to episodic memory decline call for modesty in the implications drawn.

## Acknowledgement

The Lifebrain project is funded by the EU Horizon 2020 Grant: ‘Healthy minds 0–100 years: Optimising the use of European brain imaging cohorts (“Lifebrain”)’. Grant agreement number: 732592. Call: Societal challenges: Health, demographic change and well-being. In addition, the different sub-studies are supported by different sources:m LCBC: The European Research Council under grant agreements 283634, 725025 (to A.M.F.) and 313440 (to K.B.W.), as well as the Norwegian Research Council (to A.M.F., K.B.W.), The National Association for Public Health’s dementia research program, Norway (to A.M.F) and the Medical Student Research Program at the University of Oslo. Betula: a scholar grant from the Knut and Alice Wallenberg (KAW) foundation to L.N. Barcelona: Partially supported by a Spanish Ministry of Economy and Competitiveness (MINECO) grant to D-BF [grant number PSI2015-64227-R (AEI/FEDER, UE)]; by the Walnuts and Healthy Aging study (http://www.clinicaltrials.gov; Grant NCT01634841) funded by the California Walnut Commission, Sacramento, California. BASE-II has been supported by the German Federal Ministry of Education and Research under grant numbers 16SV5537/16SV5837/16SV5538/16SV5536K/01UW0808/01UW0706/01GL1716A/01GL1716 B, and S.K. has received support from the European Research Council under grant agreement 677804.

The Wellcome Centre for Integrative Neuroimaging is supported by core funding from award 203139/Z/16/Z from the Wellcome Trust. Drs Suri and Zsoldos were funded by award the UK Medical Research Council (G1001354) and the HDH Wills 1965 Charitable Trust (1117747). Dr Suri is now funded by the UK Alzheimer’s Society Research Fellowship (Grant Ref 441); CES and Suri are supported by the NIHR Oxford Health Biomedical Research Centre. Parts of the data collection and sharing was funded by the Alzheimer’s Disease Neuroimaging Initiative (ADNI) (National Institutes of Health Grant U01 AG024904) and DOD ADNI (Department of Defense award number W81XWH-12-2-0012).

Data collection and sharing for this project was also funded by the Alzheimer’s Disease Neuroimaging Initiative (ADNI) (National Institutes of Health Grant U01 AG024904) and DOD ADNI (Department of Defense award number W81XWH-12-2-0012). ADNI is funded by the National Institute on Aging, the National Institute of Biomedical Imaging and Bioengineering, and through generous contributions from the following: AbbVie, Alzheimer’s Association; Alzheimer’s Drug Discovery Foundation; Araclon Biotech; BioClinica, Inc.; Biogen; Bristol-Myers Squibb Company; CereSpir, Inc.; Cogstate; Eisai Inc.; Elan Pharmaceuticals, Inc.; Eli Lilly and Company; EuroImmun; F. Hoffmann-La Roche Ltd and its affiliated company Genentech, Inc.; Fujirebio; GE Healthcare; IXICO Ltd.; Janssen Alzheimer Immunotherapy Research & Development, LLC.; Johnson & Johnson Pharmaceutical Research & Development LLC.; Lumosity; Lundbeck; Merck & Co., Inc.; Meso Scale Diagnostics, LLC.; NeuroRx Research; Neurotrack Technologies; Novartis Pharmaceuticals Corporation; Pfizer Inc.; Piramal Imaging; Servier; Takeda Pharmaceutical Company; and Transition Therapeutics. The Canadian Institutes of Health Research is providing funds to support ADNI clinical sites in Canada. Private sector contributions are facilitated by the Foundation for the National Institutes of Health (www.fnih.org). The grantee organization is the Northern California Institute for Research and Education, and the study is coordinated by the Alzheimer’s Therapeutic Research Institute at the University of Southern California. ADNI data are disseminated by the Laboratory for Neuro Imaging at the University of Southern California.

