## Supplementary material for "Self-reported sleep problems are related to cortical thinning in aging but not memory decline and amyloid-β accumulation – results from the Lifebrain consortium": SI

**for**

**1. Supplemental information about the samples**

**2. Conversion from KSQ to PSQI**

**3. Details on statistical analyses**

**4. Post hoc analyses controlling for BMI and depression symptoms**

**5. Details about the memory analyses**

**6. Quantification of Aβ**

**7. Virtual histology analyses**

**8. Complete listing of ADNI researchers**

**9. References**

**1. Supplemental information about the samples**

**General Descriptions**

The Lifebrain sample was derived from major European brain studies. The main features of each samples, as well as key references, are provided below.

**LCBC**

*Sample source*

Center for Lifespan Changes in Brain and Cognition

*General description of study/ procedures*

Cognitively normal participants were drawn from studies coordinated by the Research Group for Lifespan Changes in Brain and Cognition (LCBC [www.oslobrains.no](http://www.oslobrains.no)), approved by a Norwegian Regional Committee for Medical and Health Research Ethics. Written informed consent was obtained from all participants.

*Recruitment*

Newspaper adds, web page adds

*Population*

The major part of the sample (n=843) consisted of normal, cognitively healthy participants across the lifespan. One sub-population (n = 103) consisted of patients scheduled for elective gynecological (genital prolapse), urological (benign prostate hyperplasia, prostate cancer, or bladder tumor/cancer) or orthopedic (knee or hip replacement) surgery in spinal anesthesia, turning 65 years or older the year of inclusion.

*Inclusion/ exclusion criteria, screening*

Adult participants were screened using a standardized health interview prior to inclusion in the study. Participants with a history of self- or parent-reported neurological or psychiatric conditions, including clinically significant stroke, serious head injury, untreated hypertension, diabetes, and use of psychoactive drugs within the last two years, were excluded. Further, participants reporting worries concerning their cognitive status, including memory function, were excluded. All participants 40-80 years of age were required to score ≥26 and participants > 80 years ≥ 25 on the Mini Mental State Examination [1] according to population norms [2]. From the sub-population of elective surgery patients, dementia, previous stroke with sequela, Parkinson's disease, and other neurodegenerative diseases likely to affect cognitive function were initial exclusion criteria. From this pool of participants, we further selected only cognitively healthy participants based on clinical examinations at Department of Geriatric Medicine at Oslo University Hospital.

*Key references*

Langnes E, Sneve MH, Sederevicius D, Amlien IK, Walhovd KB, Fjell AM. Lifespan trajectories and relationships to memory of the macro- and microstructure of the anterior and posterior hippocampus – a longitudinal multi-modal imaging study. bioRxiv. doi: <https://doi.org/10.1101/564732>

Fjell AM, Idland AV, Sala-Llonch R, Watne LO, Borza T, Brækhus A, Lona T, Zeterberg H, Blennow K, Wyller TB, Walhovd KB. Neuroinflammation and Tau interact with amyloid in predicting sleep problems in aging independently of atrophy. Cerebral Cortex, 2018, 28, 2775-2785.

**BASE-II**

*Sample source*

Berlin Aging Study-II

*General description of study/ procedures*

The medical exam consisted of a 2-day protocol including a comprehensive anamnesis performed by a physician and involving a wide array of laboratory and functional tests (including PSQI questionnaire). Medical variables were collected about 1 year prior to cognitive testing (mean time difference in years = 1.2 years; SD = 0.80). After completion of the medical examination, participants were invited to two cognitive testing sessions scheduled 1 week apart, and were tested in small groups (e.g. about 6 participants per group) on a comprehensive cognitive battery that covers key cognitive abilities measured by 21 tasks. Each session lasted about 3.5 h. From one session to the next, participants were asked to fill out psychosocial questionnaires related to subjective health and well-being. The different elements of the study were approved by the ethics committees of the Max Planck Institute for Human Development, the Charité University ethics committee and by the ethics committees of DGPs. Participants signed written informed consent and received monetary compensation for their participation in BASE-II and the MRI study. All experiments were performed in accordance with relevant guidelines and regulations.

On average the participants had 14.01 years of education (SD = 2.89) and a body mass index of 26.70 (SD = 3.51). Most of the participants were married and still living together (63%), while 14% were divorced, 4.4% single and 4.1% widowed. None of the participants took any medication that may have affected memory function or had a history of head injuries, medical (e.g., heart attack), neurological (e.g., epilepsy), or psychiatric disorders (e.g., depression).

*Recruitment*

Baseline Sample (TP1):

Participants were community-dwelling older adults recruited from the greater Berlin

metropolitan area through advertisements in newspapers and public areas. Participants were recruited within the Berlin Aging Study II (BASE-II) (for cohort characteristics and additional details, see Bertram et al., 2014; Gerstorf et al., 2016). The baseline sample comprised 1979 participants (the original sample consists of 2200 subjects, but we reduced the sample to those of which we have sleep and/or cognitive information). Of these, 1519 were older adults aged 61–88 years (mean age 71.5, SD 3.89; 793 female), and 460 were younger adults aged 24–40years (mean age 31.1, SD 3.38; 247 female). On average, older participants had 14.59 years of education (SD 3.03), and younger participants 15.53 years (SD 2.47).

MR Sample: After completion the comprehensive cognitive examination of BASE-II, eligible participants were invited to take part in one MRI session within a time window of 2–4 weeks after cognitive testing, consisting of 341 older adults aged 61–82 years (mean age 70.1, SD = 3.89; 131 female) and 103 younger adults (mean age 31.4, SD = 3.7; 39 female).

Longitudinal data: MR scans and cognitive scores were obtained two times (a baseline (TP1): 2012-2013; and follow-up (TP2): 2015/2016). The follow-up sample (TP2) consisted of 325 participants (247 older adults, 68 younger adults) that were re-invited from the MR-subsample only. They were invited to one cognitive session, lasting 3.5 hours and another separate MR session consisting of identical measures of the baseline study.

*Population*

Community-dwelling older adults recruited from the greater Berlin metropolitan area

*Inclusion/ exclusion criteria, screening*

None of the participants took medication that might affect memory function, and none had neurological disorders, psychiatric disorders, or a history of head injuries. All participants reported normal or corrected to normal vision, were right-handed, and scored over 27 on the Mini-Mental Status Examination.

*Key references*

Bertram, L., Böckenhoff, A., Demuth, I., Düzel, S., Eckardt, R., Li, S.-C. C., … Steinhagen-Thiessen, E. (2014). Cohort profile: The Berlin Aging Study II (BASE-II). International journal of epidemiology,43(3), 703–12. doi:10.1093/ije/dyt018

Gerstorf, D., Bertram, L., Lindenberger, U., Pawelec, G., Demuth, I., Steinhagen-Thiessen, E., & Wagner, G. G. (2016). Editorial. Gerontology,62(3), 311–5. doi:10.1159/000441495

**BETULA**

*Sample source*

The Betula longitudinal study on aging, memory and dementia

*General description of study/ procedures*

A subset of 376 participants from the longitudinal Betula study (Nilsson et al., 1997) underwent structural and functional MRI in 2009-2010 and 232 returned for a follow-up scan in 2013-2014. The parent samples from which the scanned participants were derived from were originally recruited to the study in 1988, 1993, and 2008 respectively. The study is approved by the relevant ethical review board.

*Recruitment*

Population-based sampling was used for recruitment, detailed recruitment procedures are found in Nilsson et al., 1997. Participation in the neuroimaging study was offered to all participants who had remained in the study and completed cognitive testing at the 5th Betula test wave in 2008-2009.

*Population*

Population-based, healthy middle-aged and older adults

*Inclusion/ exclusion criteria, screening*

Severe visual or auditory handicaps, intellectual or developmental disabilities, suspected dementia, having a mother tongue other than Swedish, MRI contraindications, neurological disorders, or visual/motor deficits that could interfere with fMRI data collection, MMSE <24, brain or head surgery, substantial brain anatomical deviations.

*Key references*

Nilsson, L.-G., Bäckman, L., Erngrund, K., Nyberg, L., Adolfsson, R., Bucht, G., Karlsson, S., Widing, M., Winblad, B., 1997. The Betula prospective cohort study: Memory, health, and aging. Aging, Neuropsychol. Cogn. 4, 1–32. doi:10.1080/13825589708256633

**Cam-CAN**

*Sample source*

The Cambridge Centre for Ageing and Neuroscience (Cam-CAN) study

*General description of study/ procedures*

A population-based cohort of 3000 adults aged 18 was recruited to Stage 1 of the project, where they completed an interview including health and lifestyle questions, a core cognitive assessment, and a self-completed questionnaire of lifetime experiences and physical activity. Of those interviewed, ~700 participants aged 18-87 (100 per age decile) continued to Stage 2 where they undergo cognitive testing and provide measures of brain structure and function. A subset of ~250 adults returned for longitudinal follow-up data. The study is conducted in compliance with the Helsinki Declaration, and has been approved by the local ethics committee, Cambridgeshire 2 Research Ethics Committee (reference: 10/H0308/50).

*Recruitment*

Invitation letters based on the patient lists of general practitioners within the Cambridge City area

*Population*

Population-based, adult lifespan (18 years and up), cognitively healthy

*Inclusion/ exclusion criteria, screening*

General exclusion criteria: Term-time residents of colleges and universities, and participants whose Primary Care Physician feel are inappropriate to include.

Exclusion criteria for the MRI part of the study: Not cognitively normal (MMSE ≤ 24, memory defect, consent difficulties), communication difficulties (hearing problems [35db at 1000 Hz], insufficient English language, vision difficulties), medical problems by self-report of diagnosis (dementia diagnosis /Alzheimer’s Disease, Parkinson’s Disease, Motor Neurone disease, Multiple sclerosis, cancer, stroke, encephalitis, meningitis, epilepsy, head injury with serious results [coma, unconscious for >2 hours, skull fracture], recently diagnosed or uncontrolled high blood pressure, possible pregnancy, current psychiatric conditions [bipolar disorder, schizophrenia, psychosis]), mobility problems (restricted mobility which could prevent further participation, inability to walk 10 metres), substance abuse (past or current treatment for drug abuse, current drug usage), MRI/ MEG safety and comfort exclusions.

*Key references*

Shafto et al. The Cambridge Centre for Ageing and Neuroscience (Cam-CAN) study protocol: a cross-sectional, lifespan, multidisciplinary examination of healthy cognitive ageing. BMC Neurology, 2014, 14:204. doi: 10.1186/s12883-014-0204-1

**University of Barcelona**

*Sample source*

Different brain aging studies from the University of Barcelona; the WAHA cohort, CR/ iTBS cohorts, GABA cohort

*General description of study/ procedures*

Healthy middle-aged/ older adults, all gave informed consent, in accordance with the Declaration of Helsinki (1964, last revision 2013). All study procedures were approved by the local Institutional Review Board).

*Recruitment*

WAHA cohort: Eligible participants were recruited via mailing study brochures (LLU) or through the non-profit organization Institute of Aging (BCN), advertisements in the study centers, and word of mouth. Interested individuals attended an informational group meeting, completed a short medical questionnaire and signed the informed consent.

*Population*

Middle-aged and older, cognitively normal

*Inclusion/ exclusion criteria, screening*

WAHA cohort: Participants were healthy elderly men and women with normal cognitive and visual function at the time of recruitment. Inclusion criteria were age between 63 and 79 years, apparently healthy, and equally willing to be in either of the two groups. Exclusion criteria included inability to undergo neuropsychological testing; morbid obesity (BMI ≥ 40 kg/m2); uncontrolled diabetes (HbA1c > 8%); uncontrolled hypertension (on-treatment blood pressure ≥ 150/100 mmHg); prior stroke, significant head trauma or brain surgery; relevant psychiatric illness; major depression; cognitive deterioration or dementia with a score < 24 on the Mini-Mental State Examination; other neurodegenerative disorders like Parkinson’s disease; advanced AMD or eye-related conditions precluding ophthalmological evaluation; prior chemotherapy; chronic illness with projected shortened lifespan; allergy to walnuts; customary use of fish oil and/or tree nuts (> 2 servings/week) and/or other relevant sources of ALA, such as flaxseed oil or soy lecithin.

*Key references*

WAHA cohort: Rajaram S, Valls-Pedret C, Cofán M, Sabaté J, Serra-Mir M, Pérez-Heras AM, Arechiga A, Casaroli-Marano RP, Alforja S, Sala-Vila A, Doménech M, Roth I, Freitas-Simoes TM, Calvo C, López-Illamola A, Haddad E, Bitok E, Kazzi N, Huey L, Fan J, Ros E. The Walnuts and Healthy Aging Study (WAHA): Protocol for a Nutritional Intervention Trial with Walnuts on Brain Aging. Front Aging Neurosci. 2017 Jan 10;8:333.

Additional analyses were done using data obtained from the Alzheimer’s Disease Neuroimaging Initiative (ADNI) database (adni.loni.usc.edu). The ADNI was launched in 2003 as a public-private partnership, led by Principal Investigator Michael W. Weiner, MD. The primary goal of ADNI has been to test whether serial magnetic resonance imaging (MRI), positron emission tomography (PET), other biological markers, and clinical and neuropsychological assessment can be combined to measure the progression of mild cognitive impairment (MCI) and early Alzheimer’s disease (AD). For up-to-date information, see www.adni-info.org.

**2. Conversion from Karolinska Sleep Questionnaire (KSQ) to PSQI**

For Betula, Karolinska Sleep Questionnaire was used, and we therefore converted the values from the single items in KSQ to PSQI [3, 4], according to the procedures described in ^32^. In brief:

#### PSQI Component 1

PSQI Component 1 rates the answers to the question (PSQI#6)

- During the past month, how would you rate your sleep quality overall?

The options and values are given in the table below:

| Answer | Value |
| --- | --- |
| Very good | 0 |
| Fairly good | 1 |
| Fairly bad | 2 |
| Very bad | 3 |

The question in KSQ that come closest, is KSQ#5:

- Hur tycker du att du sover på det hela taget? (English: How well do you think you sleep overall?)

The options are “mycket bra”(very well), “ganska bra” (quite well), “varken bra eller dåligt” (neither well nor poorly), “ganska dåligt” (quite poorly), and “mycket dåligt” (very poorly).

The numerical scale used in Betula corresponds to the following:

| Answer | Value |
| --- | --- |
| Mycket bra | 5 |
| Ganska bra | 4 |
| Varken bra eller dåligt | 3 |
| Ganska dåligt | 2 |
| Mycket dåligt | 1 |

We used the following mapping:

| KSQ5 | PSQI_Comp1 |
| --- | --- |
| Mycket bra (5) | 0 |
| Ganska bra (4) | 1 |
| Varken bra eller dåligt (3) | 1 |
| Ganska dåligt (2) | 2 |
| Mycket dåligt (1) | 3 |

#### PSQI Component 2

PSQI component 2 measures sleep latency, and is computed from the answers to PSQI#2 (number of minutes it takes to fall asleep) and PSQI#5a (how often are you not able to fall asleep within 30 minutes). We used KSQ#9a:

- Har du haft känning av följande besvär de senaste tre månaderna? (…) Svårigheter att somna (Enligsh: Have you experienced the following troubles in the past three months? (…) Trouble falling asleep

The answers to this question are in the variable sleep_07a in the Betula dataset. The possible answers are “Aldrig” (Never), “Någon gång” (Sometimes, “Flera ggr/mån” (Several times/month), “1-2 ggr/vecka” (1-2 times/week), “3-4 ggr/vecka” (3-4 times/week), and “5 ggr eller mer /vecka” (5 times or more/week).

We used the following mapping:

| KSQ9a | PSQI_Comp2 |
| --- | --- |
| Aldrig (0) | 0 |
| Sällan (1) | 1 |
| Flera ggr/mån (2) | 1 |
| 1-2 ggr/vecka (3) | 2 |
| 3-4 ggr/vecka (4) | 3 |
| 5 ggr eller mer /vecka (5) | 3 |

#### PSQI Component 3

PSQI Component 3 ranks sleep duration (PSQI#4), using the following scale:

| Answer | Value |
| --- | --- |
| > 7 hours | 0 |
| 6-7 hours | 1 |
| 5-6 hours | 2 |
| < 5 hours | 3 |

This can be computed directly from the Betula data. The following assumptions are used:

- Columns sleep_03a_bedtime sleep_03b_bedtime contain timepoint at which the participant went to bed on weekdays and weekends, respectively.
- Columns sleep_03b_minbef and sleep_03b_minbef contain the number of minutes after bedtime until the participant was asleep on weekdays and weekends, respectively.
- Columns sleep_03a_minbef and sleep_03b_risetime contain the timepoint at which the participant woke up on weekdays and weekends, respectively.

A weighted average between weekdays (a) and weekends (b) is taken, with weights 5/7 and 2/7. If one is missing, we take the non-missing value.

#### PSQI Component 4

This component measures sleep efficiency, and can be directly calculated from the KSQ data. In order to keep this an integer, we take the weekday efficiency if it exists, otherwise the weekend efficiency.

#### PSQI Component 5

We used the following mapping between PSQI#5b-PSQI#5j and KSQ#9:

| PSQI | KSQ |
| --- | --- |
| 5b) Wake up in the middle of the night or early morning | 9c) Upprepade uppvakanden + 9i) För tidigt uppvakande (Repeated awakenings + Premature awakening) |
| 5c) Have to get up to use the bathroom | No match in KSQ. |
| 5d) Cannot breathe comfortably | 9e) Kippar efter andan, “frustar” under sömnen + 9f) Andningsuppehåll under sömnen (Gasping for breath, «snorting» during sleeping + Pauses in breathing during sleep) |
| 5e) Cough or snore loudly | 9d) Kraftiga egna snarkningar (Severe own snoring) |
| 5f) Feel too cold | No match in KSQ. |
| 5g) Feel to hot | No match in KSQ. |
| 5h) Had bad dreams | 9g) Mardrömmar. (Nightmares) |
| 5i) Have pain | No match in KSQ. |
| 5j) Other reasons | 9j) Störd/orolig sömn (Disrupted/restless sleep) |

These are a total of 7 KSQ questions, matching 9 PSQI questions. We used the same conversion as before:

| KSQ9(c-g,i-j) | PSQI_Comp2 |
| --- | --- |
| Aldrig (0) | 0 |
| Sällan (1) | 1 |
| Flera ggr/mån (2) | 1 |
| 1-2 ggr/vecka (3) | 2 |
| 3-4 ggr/vecka (4) | 3 |
| 5 ggr eller mer /vecka (5) | 3 |

The total score is summed and then multiplied by 7/9 and the component 5 score is assigned using the PSQI Component 5 calculation.

#### PSQI Component 6

No question about sleep medication is found in KSQ, so this one is missing.

#### PSQI Component 7

This component is based on PSQI#8 and PSQI#9:

**PSQI#8: During the past month, how often have you had trouble staying awake while driving, eating meals, or engaging in social activity?**

- Not during the past month (0)
- Less than once a week (1)
- Once or twice a week (2)
- Three or more times a week (3)

**PSQI#9: During the past month, how much of a problem has it been for you to keep up enough enthusiasm to get things done?**

- No problem at all (0)
- Only a very slight problem (1)
- Somewhat of a problem (2)
- A very big problem (3)

We used the following KSQ questions as proxies for PSQI Component 7:

**Har du haft känning av följande besvär de senaste tre månederna? (English: Have you experienced the following troubles during the last three months?)**

- 9m) Sömnig under arbete (Sleepiness during work)
- 9n) Sömnig under fritid (Sleepiness during spare time)
- 9o) Ofrivilliga sömnperioder (tillnickning) under arbetet (Involuntary sleep periods during work)
- 9p) Ofrivilliga sömnperioder (tillnickning) under fritid (Involuntary sleep periods durign spare time)
- 9q) Behov av att kämpa mot sömnen för att hålla sig vaken? (Need to fight sleepiness to stay awake)
- 9r) Trött i huvudet under dagen (Mental tiredness during the day)

We used the same ranking as before:

| KSQ#9q | PSQI Component 8 |
| --- | --- |
| Aldrig (0) | 0 |
| Sällan (1) | 1 |
| Flera ggr/mån (2) | 1 |
| 1-2 ggr/vecka (3) | 2 |
| 3-4 ggr/vecka (4) | 3 |
| 5 ggr eller mer /vecka (5) | 3 |

The answers are summed, and multiplied by 2/6 to get the same numerical scale as PSQI Component 7.

#### PSQI Global

This one is computed for Betula by setting PSQI Component 6 to zero, and summing the others, then multiplying by 7/6 to compensate for the lack of Component 6.

**3. Details on statistical analyses**

*Cortical surface analyses* Spatiotemporal linear mixed models^58,59^ implemented in FreeSurfer v6.0.1 and running on MATLAB R2017a were used to test relationships between sleep variables and cortical thickness/ volume change, accounting for the spatial correlation between residuals at neighboring vertices and the temporal correlation of residuals within repeated measurements of single participants. For each model, the full set of vertices was first divided into connected regions with homogeneous temporal and spatial correlations. We previously showed that power to detect sleep-*brain change* relationships are superior to power to detect sleep-*brain intercept* relationships^35^, and hence did not focus on intercept-intercept associations. Next, multivariate linear mixed models were fitted to each homogeneous region separately, assuming spatial correlations between residuals of any pair of vertices within each region to decay exponentially with Euclidean distance. Clusterwise multiple comparison correction of p-values for regression coefficients was then performed^60^, retaining clusters with a cluster-wise p-value below 0.05. Cortical thickness and volume at all vertices were separately regressed on each PSQI sleep component score, baseline age, time since baseline, site, and sex. Separate analyses included all two- and three-way interactions between PSQI, age, and time. The PSQI×time interaction represents how sleep is related to longitudinal change in thickness/volume, while the PSQI×time×age interaction represents how this relationship varies across participants’ age. The hierarchical nature of the data (repeated measurements nested within participants) was accounted for using a random intercept term. Additional analyses were conducted, in which a dichotomous indicator variable for age (greater or equal vs. lower than 60 years) interacted with PSQI scale and time, thus creating a piecewise linear model for cortical thinning with a cut-point at 60 years. All follow-up analyses were run in R^61^. To assess relationships between PSQI and memory change, we ran generalized additive mixed models^62^ (GAMM) using the package “mgcv” because they allowed us to accommodate the hypothesized non-linear effect of age on memory using smooth terms for baseline age. Again, we accounted for the nested structure with participants within studies with a random intercept. In addition, we allowed the effect of time and sleep to vary between studies, by including these as random effects per study. Additional models were run with the three-way interaction PSQI×time×baseline-age and the interaction between age at testing and sleep [s(Age,by=sleep)].

*Virtual histology* To test the relationship between sleep-related cortical thinning and cell-specific gene-expression we computed Pearson correlation coefficients between the thinning coefficients and the median inter-regional profile of gene expression levels for each marker gene, yielding a gene-specific measure of expression - thinning correlation. Using a resampling-based approach (Shin et al., 2018b), the average expression - thinning correlation for each group of cell-specific genes served as the test statistic, i.e. mean correlation between cortical phenotype and cell-specific gene expression. False discovery rate was used to account for the number of different cell-types (P_fdr_, n = 9).

*A*β*-sleep* Partial correlations were used to relate sleep to Aβ as a continuous measure from CSF (pg/ml Aβ42) or PET (PET signal in each vertex was divided by the mean signal of the cerebellum cortex to obtain standardized uptake value ratios – SUVR – see SI), controlling for age and sex, in each sample separately. The resulting coefficients were submitted to a meta-analysis (metacor, R, meta version 4.9-8). We ran additional analyses in ADNI by generalized linear mixed effects regressions (logistic link function with binominal error distribution; “lme4” R- package) since the sleep outcome in ADNI is dichotomous. Sleep disturbance was the outcome, clinical diagnosis, Aβ, age and gender were fixed effect terms and subjects were the random effects. Statistical significance was assessed with Kenward-Roger-corrected tests as implemented with the “car” R-package^63^.

**4. Post hoc analyses controlling for BMI and depression symptoms**

library(tidyverse)
library(mgcv)
library(gamm4)

In addition to visualizing the interaction effects, we investigate the effects of controling for BMI and Depression. As BMI and Depression are only available for non-overlapping subsets of participants, we have these variables in separate dataframes, and test for each variable separately.

```
prepare_data <- function(dat){
  dat %>%
    mutate(
      Site_NameAvanto = pmap_dbl(select(., starts_with("Site_Name")),
                                 function(...) as.numeric(sum(...) == 0))
    ) %>%
    pivot_longer(cols = starts_with("Site_Name")) %>%
    filter(value == 1) %>%
    mutate(
      Site_Name = case_when(
        name == "Site_NameBarcelona" ~ "Barcelona",
        name == "Site_NameBASEII" ~ "BASEII",
        name == "Site_NameBetula" ~ "Betula",
        name == "Site_NameCamCAN" ~ "CamCAN",
        name == "Site_NamePrisma" ~ "Prisma",
        name == "Site_NameSkyra" ~ "Skyra",
        name == "Site_NameAvanto" ~ "Avanto"
      )
    ) %>%
  select(ID, Site_Name, mgh_row_number) %>%
  inner_join(readRDS("data/full_data.rds"),
             by = c("ID", "Site_Name", "mgh_row_number")) %>%
  group_by(ID) %>%
  mutate(BL_Age = min(Age), Interval = Age - BL_Age) %>%
  ungroup() %>%
  select_at(vars(ID, Site, Site_Name, Age, BL_Age,
                 Interval, PSQI_Comp5_Problems, Sex, matches("BMI"),
                 matches("Depression")))
}
full_data <- read_delim("data/PSQI_Comp5_Problems.qdec.csv", delim = " ") %>%
  prepare_data()

bmi_data <- full_data %>%
  select(ID, Site_Name, Age) %>%
  left_join(
    prepare_data(read_delim("data/PSQI_Comp5_Problems.BMI.qdec.csv", delim = " ")),
    by = c("ID", "Site_Name", "Age")
    )

depression_data <- full_data %>%
  select(ID, Site_Name, Age) %>%
  left_join(
    prepare_data(read_delim("data/PSQI_Comp5_Problems.Depression.qdec.csv", delim = " ")),
    by = c("ID", "Site_Name", "Age")
    )
```

Load the different cluster thicknesses. Cluster 1 is in the left hemisphere and clusters 2-4 are in the right hemisphere.

```
Cluster1 <- read_delim("data/label_stats/20200309_lh.PSQI_Comp5_Problems.3w.psqi_time_blage.thickness.cache.th13.abs.sig_label-001.label.stats",
                       delim = ";", col_types = "c", col_names = "Average_Thickness") %>%
  mutate(Average_Thickness = as.numeric(Average_Thickness))

Cluster2 <- read_delim("data/label_stats/20200309_rh.PSQI_Comp5_Problems.3w.psqi_time_blage.thickness.cache.th13.abs.sig_label-001.label.stats",
                       delim = ";", col_types = "c", col_names = "Average_Thickness") %>%
  mutate(Average_Thickness = as.numeric(Average_Thickness))

Cluster3 <- read_delim("data/label_stats/20200309_rh.PSQI_Comp5_Problems.3w.psqi_time_blage.thickness.cache.th13.abs.sig_label-002.label.stats",
                       delim = ";", col_types = "c", col_names = "Average_Thickness") %>%
  mutate(Average_Thickness = as.numeric(Average_Thickness))

Cluster4 <- read_delim("data/label_stats/20200309_rh.PSQI_Comp5_Problems.3w.psqi_time_blage.thickness.cache.th13.abs.sig_label-003.label.stats",
                       delim = ";", col_types = "c", col_names = "Average_Thickness") %>%
  mutate(Average_Thickness = as.numeric(Average_Thickness))
```

The histograms show the distribution of BMI and Depression in the datasets for which we have it.

```
bmi_data %>%
  select(Site, BMI) %>%
  na.omit() %>%
  ggplot(aes(BMI)) +
    geom_histogram(bins = 50) +
    facet_wrap(vars(Site))
```

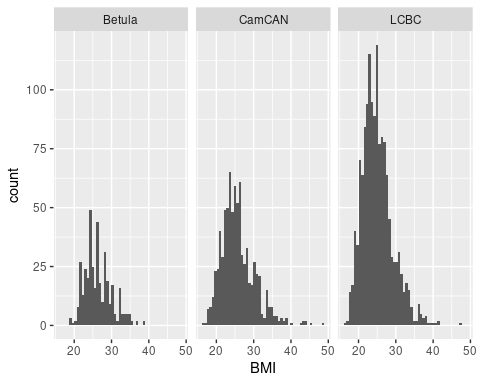

```
depression_data %>%
  select(Site, Depression) %>%
  na.omit() %>%
  ggplot(aes(Depression)) +
  geom_histogram(bins = 20) +
  facet_wrap(vars(Site))
```

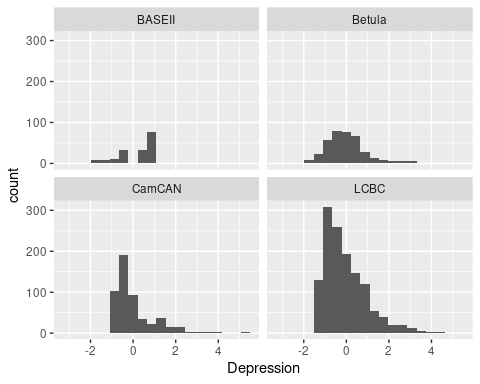

### Clusters for Age x Time x PSQI_Component5

#### Cluster 1

First we show some descriptive plots in the cluster.

```
dat1 <- full_data %>%
  bind_cols(Cluster1)

dat1_bmi <- bmi_data %>%
  bind_cols(Cluster1)

dat1_depression <- depression_data %>%
  bind_cols(Cluster1)
```

Thickness decreases with age.

```
ggplot(dat1, aes(x = Age, y = Average_Thickness)) +
  geom_line(aes(group = ID), alpha = .2) +
  geom_point(alpha = .2) +
  geom_smooth(method = "lm", se = FALSE) +
  theme_classic()
```

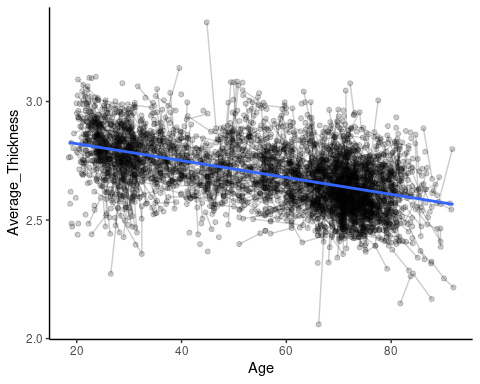

The relationship between PSQI Component 5 and thickness is very modest.

```
ggplot(dat1, aes(x = PSQI_Comp5_Problems, y = Average_Thickness)) +
  geom_line(aes(group = ID), alpha = .2) +
  geom_point(alpha = .2) +
  geom_smooth(method = "lm", se = FALSE) +
  theme_classic()
```

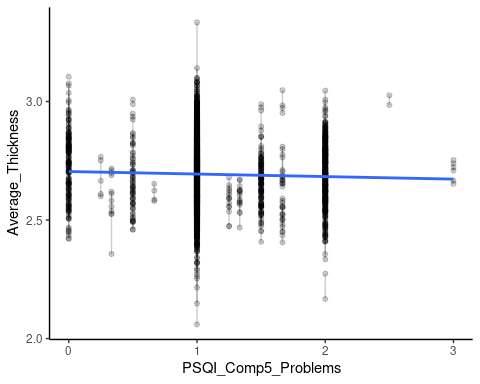

We now fit a GAMM with an interaction between Age and PSQI to visualize the relationships. The formulation below defines a smooth function of age and a linear regression term for PSQI_Comp5_Problems whose slope depends on age. Because the PSQI variable only takes on a discrete set of four values, we do not define a full smooth term for this one.

```
mod1 <- gamm4(Average_Thickness ~ s(Age) + s(Age, by = PSQI_Comp5_Problems, k = 6) + Sex + Site_Name,
              data = dat1, random = ~ (1|ID))
```

We set up a grid over which to plot the fits, and then compute and plot.

```
grid <- crossing(
  Site_Name = "Avanto",
  Sex = "Female",
  BL_Age = seq(from = 20, to = 90, by = 10),
  Interval = seq(from = 0, to = 5, by = 1),
  PSQI_Comp5_Problems = 0:3
) %>%
  mutate(Age = BL_Age + Interval)

prediction <- predict(mod1$gam, newdata = grid, se.fit = TRUE)

preds <- grid %>%
  mutate(
    fit = as.numeric(prediction$fit),
    se.fit = as.numeric(prediction$se.fit),
    BL_Age = factor(BL_Age),
    PSQI_Comp5_Problems = factor(PSQI_Comp5_Problems)
  )

p1 <- ggplot(preds, aes(x = Interval, y = fit,
                        ymin = fit + qnorm(.025) * se.fit,
                        ymax = fit + qnorm(.975) * se.fit,
                        color = PSQI_Comp5_Problems,
                        fill = PSQI_Comp5_Problems
                        )) +
  geom_line() +
  geom_ribbon(alpha = .2, color = NA) +
  facet_wrap(vars(BL_Age), scales = "free_y",
             labeller = as_labeller(function(x) paste("BL_Age =", x))) +
  theme_classic() +
  theme(legend.position = "bottom")

p1
```

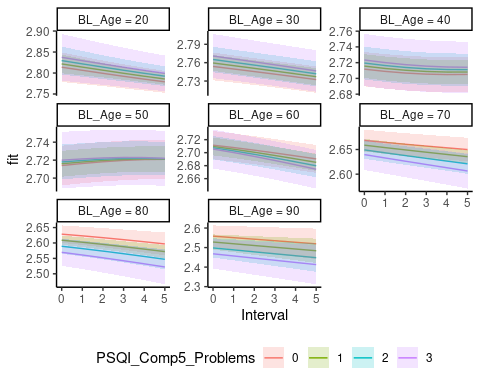

```
p2 <- ggplot(preds, aes(x = Interval, y = fit,
                        group = PSQI_Comp5_Problems,
                        color = PSQI_Comp5_Problems)) +
  geom_line() +
  facet_wrap(vars(BL_Age),
             labeller = as_labeller(function(x) paste("BL_Age =", x))) +
  theme_classic() +
  theme(legend.position = "bottom")

p2
```

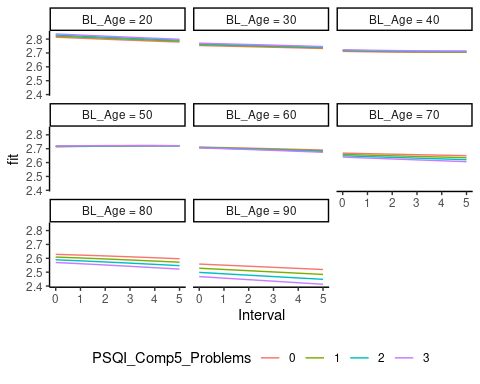

```
## Saving 5 x 4 in image
## Saving 5 x 4 in image
```

#### Controling for BMI

We next run to new models controling for BMI. Because the missingness of BMI might depend on other person characteristics, and definitely depends on cohort, we start by fitting the GAMM without controling for BMI on the dataset containing subjects with BMI values.

```
mod1_bmi_subjects <- gamm4(Average_Thickness ~ s(Age, k = 5) + s(Age, by = PSQI_Comp5_Problems, k = 5) +
                             Sex + Site_Name, data = dat1_bmi, random = ~(1|ID))
```

Next, we control for BMI by adding this as a linear predictor.

```
mod1_bmi_subjects_controled <- gamm4(Average_Thickness ~ s(Age, k = 5) + s(Age, by = PSQI_Comp5_Problems, k = 5) +
                             Sex + Site_Name + BMI, data = dat1_bmi, random = ~(1|ID))

summary(mod1_bmi_subjects_controled$gam)
##
## Family: gaussian
## Link function: identity
##
## Formula:
## Average_Thickness ~ s(Age, k = 5) + s(Age, by = PSQI_Comp5_Problems,
##     k = 5) + Sex + Site_Name + BMI
##
## Parametric coefficients:
##                   Estimate Std. Error t value Pr(>|t|)
## (Intercept)      2.6769412  0.0175394 152.625  < 2e-16 ***
## SexMale         -0.0078610  0.0053220  -1.477 0.139782
## Site_NameBetula -0.0653672  0.0086898  -7.522  7.4e-14 ***
## Site_NameCamCAN  0.0688553  0.0071578   9.620  < 2e-16 ***
## Site_NamePrisma  0.0427078  0.0126921   3.365 0.000777 ***
## Site_NameSkyra   0.0171605  0.0054160   3.168 0.001550 **
## BMI              0.0009926  0.0006160   1.611 0.107195
## ---
## Signif. codes:  0 '***' 0.001 '**' 0.01 '*' 0.05 '.' 0.1 ' ' 1
##
## Approximate significance of smooth terms:
##                              edf Ref.df      F  p-value
## s(Age)                     3.766  3.766 23.041 3.83e-14 ***
## s(Age):PSQI_Comp5_Problems 2.181  2.181  2.006    0.127
## ---
## Signif. codes:  0 '***' 0.001 '**' 0.01 '*' 0.05 '.' 0.1 ' ' 1
##
## R-sq.(adj) =  0.339
## lmer.REML = -4444.4  Scale est. = 0.0042998  n = 2578
```

From the model output, the effects look very similar. We check this by plotting, and see that the fits are indistinguishable with and without controling for BMI.

```
grid <- crossing(
  Site_Name = "Avanto",
  Sex = "Female",
  BL_Age = seq(from = 20, to = 90, by = 10),
  Interval = seq(from = 0, to = 5, by = 1),
  PSQI_Comp5_Problems = 0:3, BMI = 20
) %>%
  mutate(Age = BL_Age + Interval)

prediction_bmi_subjects <- predict(mod1_bmi_subjects$gam, newdata = grid,
                                   type = "terms",
                                   terms = c("s(Age)", "s(Age):PSQI_Comp5_Problems")) %>%
  rowSums()
prediction_bmi_subjects_controled <- predict(mod1_bmi_subjects_controled$gam, newdata = grid,
                                   type = "terms",
                                   terms = c("s(Age)", "s(Age):PSQI_Comp5_Problems")) %>%
  rowSums()

preds <- grid %>%
  mutate(
    `without BMI` = prediction_bmi_subjects,
    `with BMI` = prediction_bmi_subjects_controled,
    BL_Age = factor(BL_Age),
    PSQI_Comp5_Problems = factor(PSQI_Comp5_Problems)
  ) %>%
  pivot_longer(cols = c("without BMI", "with BMI"))

ggplot(preds, aes(x = Interval, y = value,
                  group = paste(PSQI_Comp5_Problems, name),
                  color = PSQI_Comp5_Problems,
                  linetype = name)) +
  geom_line() +
  facet_wrap(vars(BL_Age), scales = "free_y",
             labeller = as_labeller(function(x) paste("BL_Age =", x))) +
  theme_classic() +
  theme(legend.position = "bottom") +
  labs(linetype = NULL)
```

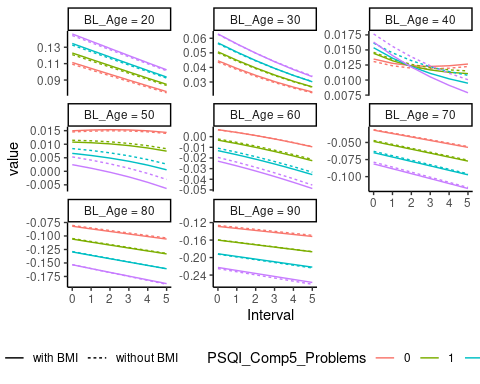

#### Controling for Depression

We next run to new models controling for Depression. Because the missingness of Depression might depend on other person characteristics, and definitely depends on cohort, we start by fitting the GAMM without controling for Depression on the dataset containing subjects with Depression values.

```
mod1_depression_subjects <- gamm4(Average_Thickness ~ s(Age, k = 5) + s(Age, by = PSQI_Comp5_Problems, k = 5) +
                             Sex + Site_Name, data = dat1_depression, random = ~(1|ID))
```

Next, we control for Depression by adding this as a linear predictor.

```
mod1_depression_subjects_controled <- gamm4(Average_Thickness ~ s(Age, k = 5) + s(Age, by = PSQI_Comp5_Problems, k = 5) +
                             Sex + Site_Name + Depression, data = dat1_depression, random = ~(1|ID))
```

From the model output, the effects look very similar. We check this by plotting, and see that the fits are indistinguishable with and without controling for Depression.

```
grid <- crossing(
  Site_Name = "Avanto",
  Sex = "Female",
  BL_Age = seq(from = 20, to = 90, by = 10),
  Interval = seq(from = 0, to = 5, by = 1),
  PSQI_Comp5_Problems = 0:3, Depression = 0
) %>%
  mutate(Age = BL_Age + Interval)

prediction_depression_subjects <- predict(mod1_depression_subjects$gam, newdata = grid,
                                   type = "terms",
                                   terms = c("s(Age)", "s(Age):PSQI_Comp5_Problems")) %>%
  rowSums()
prediction_depression_subjects_controled <- predict(mod1_depression_subjects_controled$gam, newdata = grid,
                                   type = "terms",
                                   terms = c("s(Age)", "s(Age):PSQI_Comp5_Problems")) %>%
  rowSums()

preds <- grid %>%
  mutate(
    `without Depression` = prediction_depression_subjects,
    `with Depression` = prediction_depression_subjects_controled,
    BL_Age = factor(BL_Age),
    PSQI_Comp5_Problems = factor(PSQI_Comp5_Problems)
  ) %>%
  pivot_longer(cols = c("without Depression", "with Depression"))

ggplot(preds, aes(x = Interval, y = value,
                  group = paste(PSQI_Comp5_Problems, name),
                  color = PSQI_Comp5_Problems,
                  linetype = name)) +
  geom_line() +
  facet_wrap(vars(BL_Age), scales = "free_y",
             labeller = as_labeller(function(x) paste("BL_Age =", x))) +
  theme_classic() +
  theme(legend.position = "bottom") +
  labs(linetype = NULL)
```

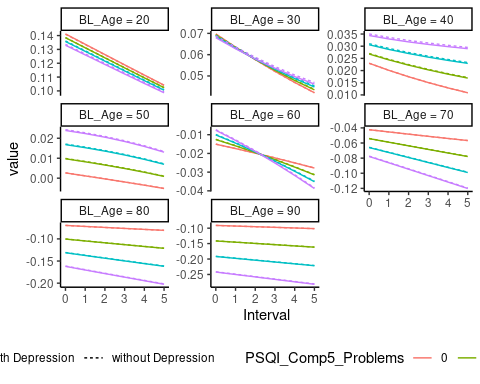

```
rm(dat1_bmi, dat1_depression, mod1_bmi_subjects,
   mod1_bmi_subjects_controled, mod1_depression_subjects,
   mod1_depression_subjects_controled)
```

### Cluster 2

First we show some descriptive plots in the cluster.

```
dat2 <- full_data %>%
  bind_cols(Cluster2)

dat2_bmi <- bmi_data %>%
  bind_cols(Cluster2)

dat2_depression <- depression_data %>%
  bind_cols(Cluster2)
```

Thickness decreases with age.

```
ggplot(dat2, aes(x = Age, y = Average_Thickness)) +
  geom_line(aes(group = ID), alpha = .2) +
  geom_point(alpha = .2) +
  geom_smooth(method = "lm", se = FALSE) +
  theme_classic()
```

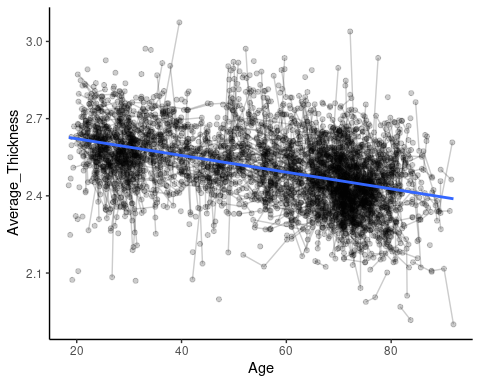

The relationship between PSQI Component 5 and thickness is very modest.

```
ggplot(dat2, aes(x = PSQI_Comp5_Problems, y = Average_Thickness)) +
  geom_line(aes(group = ID), alpha = .2) +
  geom_point(alpha = .2) +
  geom_smooth(method = "lm", se = FALSE) +
  theme_classic()
```

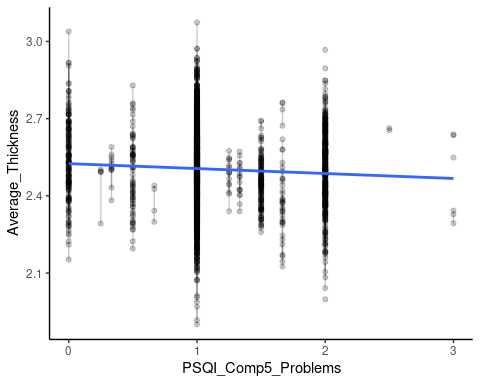

We now fit a GAMM with an interaction between Age and PSQI to visualize the relationships. The formulation below defines a smooth function of age and a linear regression term for PSQI_Comp5_Problems whose slope depends on age. Because the PSQI variable only takes on a discrete set of four values, we do not define a full smooth term for this one.

```
mod2 <- gamm4(Average_Thickness ~ s(Age) + s(Age, by = PSQI_Comp5_Problems, k = 6) + Sex + Site_Name,
              data = dat2, random = ~ (1|ID))
```

We set up a grid over which to plot the fits, and then compute and plot.

```
grid <- crossing(
  Site_Name = "Avanto",
  Sex = "Female",
  BL_Age = seq(from = 20, to = 90, by = 10),
  Interval = seq(from = 0, to = 5, by = 1),
  PSQI_Comp5_Problems = 0:3
) %>%
  mutate(Age = BL_Age + Interval)

prediction <- predict(mod2$gam, newdata = grid, se.fit = TRUE)

preds <- grid %>%
  mutate(
    fit = as.numeric(prediction$fit),
    se.fit = as.numeric(prediction$se.fit),
    BL_Age = factor(BL_Age),
    PSQI_Comp5_Problems = factor(PSQI_Comp5_Problems)
  )

p1 <- ggplot(preds, aes(x = Interval, y = fit,
                        ymin = fit + qnorm(.025) * se.fit,
                        ymax = fit + qnorm(.975) * se.fit,
                        color = PSQI_Comp5_Problems,
                        fill = PSQI_Comp5_Problems
                        )) +
  geom_line() +
  geom_ribbon(alpha = .2, color = NA) +
  facet_wrap(vars(BL_Age), scales = "free_y",
             labeller = as_labeller(function(x) paste("BL_Age =", x))) +
  theme_classic() +
  theme(legend.position = "bottom")

p1
```

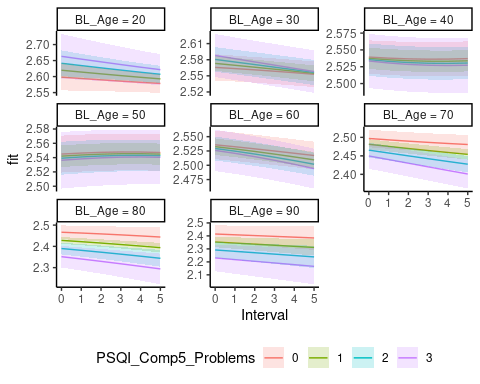

```
p2 <- ggplot(preds, aes(x = Interval, y = fit,
                        group = PSQI_Comp5_Problems,
                        color = PSQI_Comp5_Problems)) +
  geom_line() +
  facet_wrap(vars(BL_Age),
             labeller = as_labeller(function(x) paste("BL_Age =", x))) +
  theme_classic() +
  theme(legend.position = "bottom")

p2
```

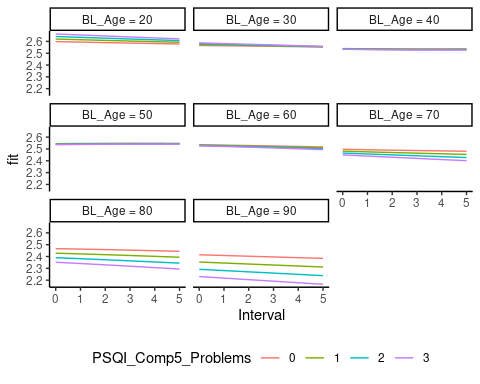

```
## Saving 5 x 4 in image
## Saving 5 x 4 in image
```

#### Controling for BMI

We next run to new models controling for BMI. Because the missingness of BMI might depend on other person characteristics, and definitely depends on cohort, we start by fitting the GAMM without controling for BMI on the dataset containing subjects with BMI values.

```
mod2_bmi_subjects <- gamm4(Average_Thickness ~ s(Age, k = 5) + s(Age, by = PSQI_Comp5_Problems, k = 5) +
                             Sex + Site_Name, data = dat2_bmi, random = ~(1|ID))
```

Next, we control for BMI by adding this as a linear predictor.

```
mod2_bmi_subjects_controled <- gamm4(Average_Thickness ~ s(Age, k = 5) + s(Age, by = PSQI_Comp5_Problems, k = 5) +
                             Sex + Site_Name + BMI, data = dat2_bmi, random = ~(1|ID))

summary(mod2_bmi_subjects_controled$gam)
##
## Family: gaussian
## Link function: identity
##
## Formula:
## Average_Thickness ~ s(Age, k = 5) + s(Age, by = PSQI_Comp5_Problems,
##     k = 5) + Sex + Site_Name + BMI
##
## Parametric coefficients:
##                   Estimate Std. Error t value Pr(>|t|)
## (Intercept)      2.4568034  0.0189171 129.872  < 2e-16 ***
## SexMale         -0.0155061  0.0056983  -2.721  0.00655 **
## Site_NameBetula -0.0786903  0.0093761  -8.393  < 2e-16 ***
## Site_NameCamCAN  0.0846573  0.0077668  10.900  < 2e-16 ***
## Site_NamePrisma  0.0711432  0.0137879   5.160 2.66e-07 ***
## Site_NameSkyra  -0.0037041  0.0061434  -0.603  0.54660
## BMI              0.0026126  0.0006636   3.937 8.47e-05 ***
## ---
## Signif. codes:  0 '***' 0.001 '**' 0.01 '*' 0.05 '.' 0.1 ' ' 1
##
## Approximate significance of smooth terms:
##                              edf Ref.df      F  p-value
## s(Age)                     1.455  1.455 17.427 1.08e-05 ***
## s(Age):PSQI_Comp5_Problems 4.513  4.513  6.222 0.000378 ***
## ---
## Signif. codes:  0 '***' 0.001 '**' 0.01 '*' 0.05 '.' 0.1 ' ' 1
##
## R-sq.(adj) =  0.322
## lmer.REML = -3950.6  Scale est. = 0.0057968  n = 2578
```

From the model output, the effects look very similar. We check this by plotting, and see that the fits are indistinguishable with and without controling for BMI.

```
grid <- crossing(
  Site_Name = "Avanto",
  Sex = "Female",
  BL_Age = seq(from = 20, to = 90, by = 10),
  Interval = seq(from = 0, to = 5, by = 1),
  PSQI_Comp5_Problems = 0:3, BMI = 20
) %>%
  mutate(Age = BL_Age + Interval)

prediction_bmi_subjects <- predict(mod2_bmi_subjects$gam, newdata = grid,
                                   type = "terms",
                                   terms = c("s(Age)", "s(Age):PSQI_Comp5_Problems")) %>%
  rowSums()
prediction_bmi_subjects_controled <- predict(mod2_bmi_subjects_controled$gam, newdata = grid,
                                   type = "terms",
                                   terms = c("s(Age)", "s(Age):PSQI_Comp5_Problems")) %>%
  rowSums()

preds <- grid %>%
  mutate(
    `without BMI` = prediction_bmi_subjects,
    `with BMI` = prediction_bmi_subjects_controled,
    BL_Age = factor(BL_Age),
    PSQI_Comp5_Problems = factor(PSQI_Comp5_Problems)
  ) %>%
  pivot_longer(cols = c("without BMI", "with BMI"))

ggplot(preds, aes(x = Interval, y = value,
                  group = paste(PSQI_Comp5_Problems, name),
                  color = PSQI_Comp5_Problems,
                  linetype = name)) +
  geom_line() +
  facet_wrap(vars(BL_Age), scales = "free_y",
             labeller = as_labeller(function(x) paste("BL_Age =", x))) +
  theme_classic() +
  theme(legend.position = "bottom") +
  labs(linetype = NULL)
```

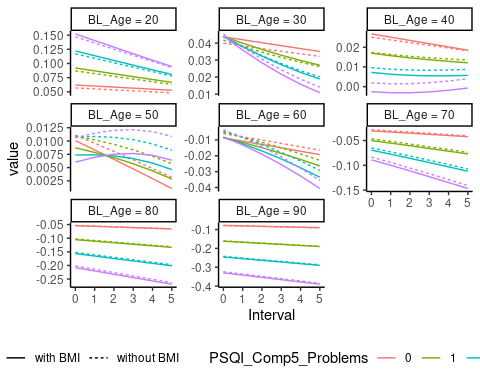

```
ggsave("/tsd/p274/data/durable/file-export/cluster2_bmi.png", dpi = 600,
       height = 12, width = 16, units = "cm")
```

#### Controling for Depression

We next run to new models controling for Depression. Because the missingness of Depression might depend on other person characteristics, and definitely depends on cohort, we start by fitting the GAMM without controling for Depression on the dataset containing subjects with Depression values.

```
mod2_depression_subjects <- gamm4(Average_Thickness ~ s(Age, k = 5) + s(Age, by = PSQI_Comp5_Problems, k = 5) +
                             Sex + Site_Name, data = dat2_depression, random = ~(1|ID))
```

Next, we control for Depression by adding this as a linear predictor.

```
mod2_depression_subjects_controled <- gamm4(Average_Thickness ~ s(Age, k = 5) + s(Age, by = PSQI_Comp5_Problems, k = 5) +
                             Sex + Site_Name + Depression, data = dat2_depression, random = ~(1|ID))
```

From the model output, the effects look very similar. We check this by plotting, and see that the fits are indistinguishable with and without controling for Depression.

```
grid <- crossing(
  Site_Name = "Avanto",
  Sex = "Female",
  BL_Age = seq(from = 20, to = 90, by = 10),
  Interval = seq(from = 0, to = 5, by = 1),
  PSQI_Comp5_Problems = 0:3, Depression = 0
) %>%
  mutate(Age = BL_Age + Interval)

prediction_depression_subjects <- predict(mod2_depression_subjects$gam, newdata = grid,
                                   type = "terms",
                                   terms = c("s(Age)", "s(Age):PSQI_Comp5_Problems")) %>%
  rowSums()
prediction_depression_subjects_controled <- predict(mod2_depression_subjects_controled$gam, newdata = grid,
                                   type = "terms",
                                   terms = c("s(Age)", "s(Age):PSQI_Comp5_Problems")) %>%
  rowSums()

preds <- grid %>%
  mutate(
    `without Depression` = prediction_depression_subjects,
    `with Depression` = prediction_depression_subjects_controled,
    BL_Age = factor(BL_Age),
    PSQI_Comp5_Problems = factor(PSQI_Comp5_Problems)
  ) %>%
  pivot_longer(cols = c("without Depression", "with Depression"))

ggplot(preds, aes(x = Interval, y = value,
                  group = paste(PSQI_Comp5_Problems, name),
                  color = PSQI_Comp5_Problems,
                  linetype = name)) +
  geom_line() +
  facet_wrap(vars(BL_Age), scales = "free_y",
             labeller = as_labeller(function(x) paste("BL_Age =", x))) +
  theme_classic() +
  theme(legend.position = "bottom") +
  labs(linetype = NULL)
```

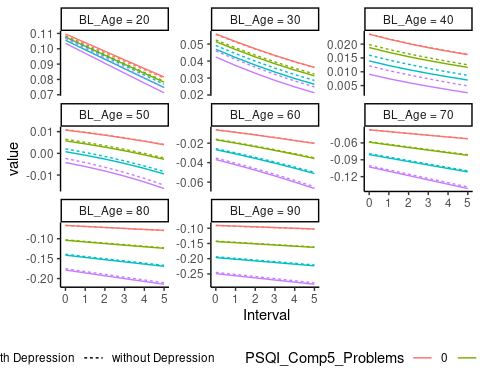

```
ggsave("/tsd/p274/data/durable/file-export/cluster2_depression.png",
       height = 12, width = 16, units = "cm", dpi = 600)
rm(dat2_bmi, dat2_depression, mod2_bmi_subjects,
   mod2_bmi_subjects_controled, mod2_depression_subjects,
   mod2_depression_subjects_controled)
```

### Cluster 3

First we show some descriptive plots in the cluster.

```
dat3 <- full_data %>%
  bind_cols(Cluster3)

dat3_bmi <- bmi_data %>%
  bind_cols(Cluster3)

dat3_depression <- depression_data %>%
  bind_cols(Cluster3)
```

Thickness decreases with age.

```
ggplot(dat3, aes(x = Age, y = Average_Thickness)) +
  geom_line(aes(group = ID), alpha = .2) +
  geom_point(alpha = .2) +
  geom_smooth(method = "lm", se = FALSE) +
  theme_classic()
```

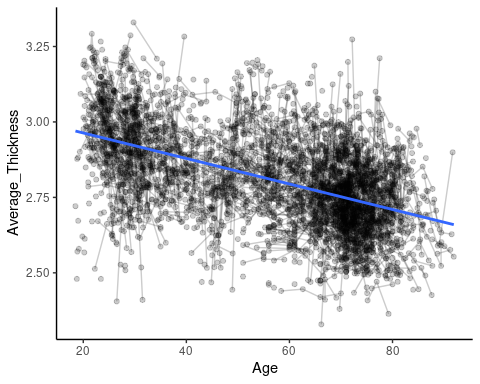

The relationship between PSQI Component 5 and thickness is very modest.

```
ggplot(dat3, aes(x = PSQI_Comp5_Problems, y = Average_Thickness)) +
  geom_line(aes(group = ID), alpha = .2) +
  geom_point(alpha = .2) +
  geom_smooth(method = "lm", se = FALSE) +
  theme_classic()
```

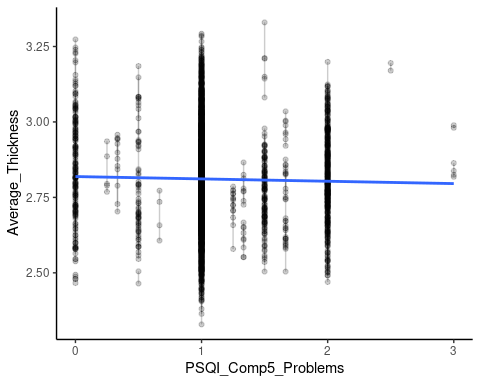

We now fit a GAMM with an interaction between Age and PSQI to visualize the relationships. The formulation below defines a smooth function of age and a linear regression term for PSQI_Comp5_Problems whose slope depends on age. Because the PSQI variable only takes on a discrete set of four values, we do not define a full smooth term for this one.

```
mod3 <- gamm4(Average_Thickness ~ s(Age) + s(Age, by = PSQI_Comp5_Problems, k = 6) + Sex + Site_Name,
              data = dat3, random = ~ (1|ID))
```

We set up a grid over which to plot the fits, and then compute and plot.

```
grid <- crossing(
  Site_Name = "Avanto",
  Sex = "Female",
  BL_Age = seq(from = 20, to = 90, by = 10),
  Interval = seq(from = 0, to = 5, by = 1),
  PSQI_Comp5_Problems = 0:3
) %>%
  mutate(Age = BL_Age + Interval)

prediction <- predict(mod3$gam, newdata = grid, se.fit = TRUE)

preds <- grid %>%
  mutate(
    fit = as.numeric(prediction$fit),
    se.fit = as.numeric(prediction$se.fit),
    BL_Age = factor(BL_Age),
    PSQI_Comp5_Problems = factor(PSQI_Comp5_Problems)
  )

p1 <- ggplot(preds, aes(x = Interval, y = fit,
                        ymin = fit + qnorm(.025) * se.fit,
                        ymax = fit + qnorm(.975) * se.fit,
                        color = PSQI_Comp5_Problems,
                        fill = PSQI_Comp5_Problems
                        )) +
  geom_line() +
  geom_ribbon(alpha = .2, color = NA) +
  facet_wrap(vars(BL_Age), scales = "free_y",
             labeller = as_labeller(function(x) paste("BL_Age =", x))) +
  theme_classic() +
  theme(legend.position = "bottom")

p1
```

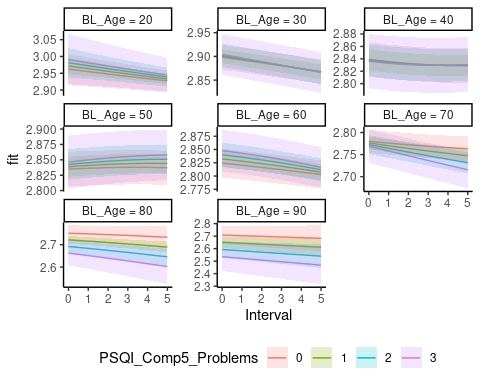

```
p2 <- ggplot(preds, aes(x = Interval, y = fit,
                        group = PSQI_Comp5_Problems,
                        color = PSQI_Comp5_Problems)) +
  geom_line() +
  facet_wrap(vars(BL_Age),
             labeller = as_labeller(function(x) paste("BL_Age =", x))) +
  theme_classic() +
  theme(legend.position = "bottom")

p2
```

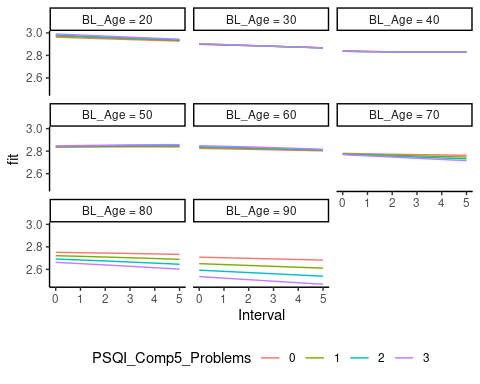

```
## Saving 5 x 4 in image
## Saving 5 x 4 in image
```

#### Controling for BMI

We next run to new models controling for BMI. Because the missingness of BMI might depend on other person characteristics, and definitely depends on cohort, we start by fitting the GAMM without controling for BMI on the dataset containing subjects with BMI values.

```
mod3_bmi_subjects <- gamm4(Average_Thickness ~ s(Age, k = 5) + s(Age, by = PSQI_Comp5_Problems, k = 5) +
                             Sex + Site_Name, data = dat3_bmi, random = ~(1|ID))
```

Next, we control for BMI by adding this as a linear predictor.

```
mod3_bmi_subjects_controled <- gamm4(Average_Thickness ~ s(Age, k = 5) + s(Age, by = PSQI_Comp5_Problems, k = 5) +
                             Sex + Site_Name + BMI, data = dat3_bmi, random = ~(1|ID))

summary(mod3_bmi_subjects_controled$gam)
##
## Family: gaussian
## Link function: identity
##
## Formula:
## Average_Thickness ~ s(Age, k = 5) + s(Age, by = PSQI_Comp5_Problems,
##     k = 5) + Sex + Site_Name + BMI
##
## Parametric coefficients:
##                   Estimate Std. Error t value Pr(>|t|)
## (Intercept)      2.764e+00  2.002e-02 138.036  < 2e-16 ***
## SexMale         -2.194e-05  6.062e-03  -0.004 0.997112
## Site_NameBetula -1.210e-01  9.926e-03 -12.190  < 2e-16 ***
## Site_NameCamCAN  3.769e-02  8.191e-03   4.602  4.4e-06 ***
## Site_NamePrisma  5.216e-02  1.452e-02   3.593 0.000333 ***
## Site_NameSkyra   2.296e-02  6.281e-03   3.655 0.000262 ***
## BMI              2.311e-03  7.031e-04   3.287 0.001027 **
## ---
## Signif. codes:  0 '***' 0.001 '**' 0.01 '*' 0.05 '.' 0.1 ' ' 1
##
## Approximate significance of smooth terms:
##                             edf Ref.df      F  p-value
## s(Age)                     3.77   3.77 26.153 2.42e-16 ***
## s(Age):PSQI_Comp5_Problems 2.00   2.00  0.752    0.472
## ---
## Signif. codes:  0 '***' 0.001 '**' 0.01 '*' 0.05 '.' 0.1 ' ' 1
##
## R-sq.(adj) =  0.357
## lmer.REML = -3731.8  Scale est. = 0.005863  n = 2578
```

From the model output, the effects look very similar. We check this by plotting, and see that the fits are indistinguishable with and without controling for BMI.

```
grid <- crossing(
  Site_Name = "Avanto",
  Sex = "Female",
  BL_Age = seq(from = 20, to = 90, by = 10),
  Interval = seq(from = 0, to = 5, by = 1),
  PSQI_Comp5_Problems = 0:3, BMI = 20
) %>%
  mutate(Age = BL_Age + Interval)

prediction_bmi_subjects <- predict(mod3_bmi_subjects$gam, newdata = grid,
                                   type = "terms",
                                   terms = c("s(Age)", "s(Age):PSQI_Comp5_Problems")) %>%
  rowSums()
prediction_bmi_subjects_controled <- predict(mod3_bmi_subjects_controled$gam, newdata = grid,
                                   type = "terms",
                                   terms = c("s(Age)", "s(Age):PSQI_Comp5_Problems")) %>%
  rowSums()

preds <- grid %>%
  mutate(
    `without BMI` = prediction_bmi_subjects,
    `with BMI` = prediction_bmi_subjects_controled,
    BL_Age = factor(BL_Age),
    PSQI_Comp5_Problems = factor(PSQI_Comp5_Problems)
  ) %>%
  pivot_longer(cols = c("without BMI", "with BMI"))

ggplot(preds, aes(x = Interval, y = value,
                  group = paste(PSQI_Comp5_Problems, name),
                  color = PSQI_Comp5_Problems,
                  linetype = name)) +
  geom_line() +
  facet_wrap(vars(BL_Age), scales = "free_y",
             labeller = as_labeller(function(x) paste("BL_Age =", x))) +
  theme_classic() +
  theme(legend.position = "bottom") +
  labs(linetype = NULL)
```

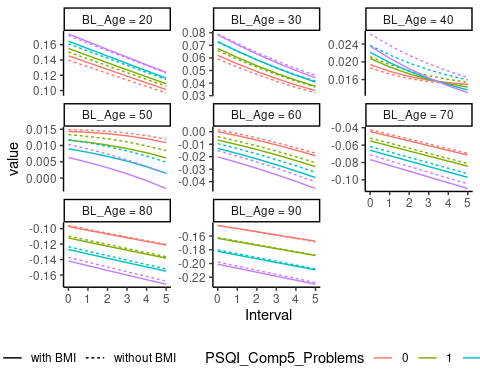

#### Controling for Depression

We next run to new models controling for Depression. Because the missingness of Depression might depend on other person characteristics, and definitely depends on cohort, we start by fitting the GAMM without controling for Depression on the dataset containing subjects with Depression values.

```
mod3_depression_subjects <- gamm4(Average_Thickness ~ s(Age, k = 5) + s(Age, by = PSQI_Comp5_Problems, k = 5) +
                             Sex + Site_Name, data = dat3_depression, random = ~(1|ID))
```

Next, we control for Depression by adding this as a linear predictor.

```
mod3_depression_subjects_controled <- gamm4(Average_Thickness ~ s(Age, k = 5) + s(Age, by = PSQI_Comp5_Problems, k = 5) +
                             Sex + Site_Name + Depression, data = dat3_depression, random = ~(1|ID))
```

From the model output, the effects look very similar. We check this by plotting, and see that the fits are indistinguishable with and without controling for Depression.

```
grid <- crossing(
  Site_Name = "Avanto",
  Sex = "Female",
  BL_Age = seq(from = 20, to = 90, by = 10),
  Interval = seq(from = 0, to = 5, by = 1),
  PSQI_Comp5_Problems = 0:3, Depression = 0
) %>%
  mutate(Age = BL_Age + Interval)

prediction_depression_subjects <- predict(mod3_depression_subjects$gam, newdata = grid,
                                   type = "terms",
                                   terms = c("s(Age)", "s(Age):PSQI_Comp5_Problems")) %>%
  rowSums()
prediction_depression_subjects_controled <- predict(mod3_depression_subjects_controled$gam, newdata = grid,
                                   type = "terms",
                                   terms = c("s(Age)", "s(Age):PSQI_Comp5_Problems")) %>%
  rowSums()

preds <- grid %>%
  mutate(
    `without Depression` = prediction_depression_subjects,
    `with Depression` = prediction_depression_subjects_controled,
    BL_Age = factor(BL_Age),
    PSQI_Comp5_Problems = factor(PSQI_Comp5_Problems)
  ) %>%
  pivot_longer(cols = c("without Depression", "with Depression"))

ggplot(preds, aes(x = Interval, y = value,
                  group = paste(PSQI_Comp5_Problems, name),
                  color = PSQI_Comp5_Problems,
                  linetype = name)) +
  geom_line() +
  facet_wrap(vars(BL_Age), scales = "free_y",
             labeller = as_labeller(function(x) paste("BL_Age =", x))) +
  theme_classic() +
  theme(legend.position = "bottom") +
  labs(linetype = NULL)
```

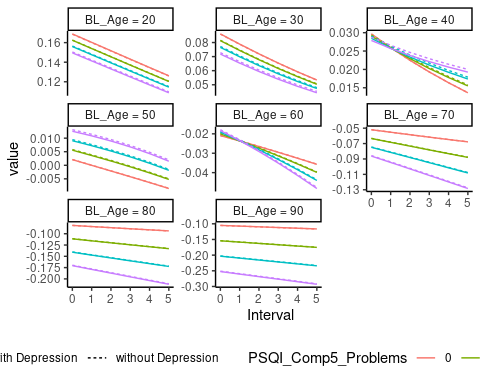

```
rm(dat3_bmi, dat3_depression, mod3_bmi_subjects,
   mod3_bmi_subjects_controled, mod3_depression_subjects,
   mod3_depression_subjects_controled)
```

### Cluster 4

First we show some descriptive plots in the cluster.

```
dat4 <- full_data %>%
  bind_cols(Cluster4)

dat4_bmi <- bmi_data %>%
  bind_cols(Cluster4)

dat4_depression <- depression_data %>%
  bind_cols(Cluster4)
```

Thickness decreases with age.

```
ggplot(dat4, aes(x = Age, y = Average_Thickness)) +
  geom_line(aes(group = ID), alpha = .2) +
  geom_point(alpha = .2) +
  geom_smooth(method = "lm", se = FALSE) +
  theme_classic()
```

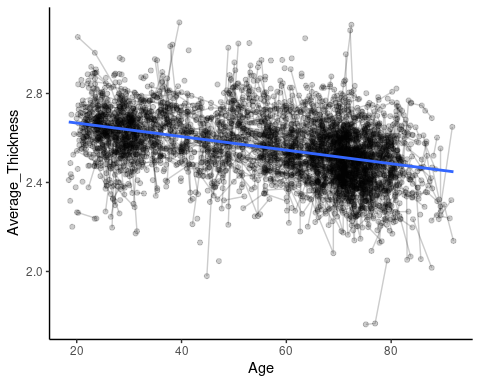

The relationship between PSQI Component 5 and thickness is very modest.

```
ggplot(dat4, aes(x = PSQI_Comp5_Problems, y = Average_Thickness)) +
  geom_line(aes(group = ID), alpha = .2) +
  geom_point(alpha = .2) +
  geom_smooth(method = "lm", se = FALSE) +
  theme_classic()
```

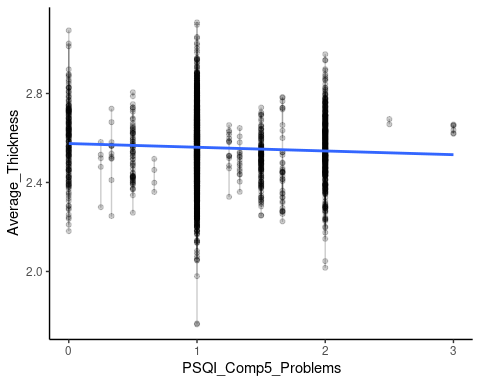

We now fit a GAMM with an interaction between Age and PSQI to visualize the relationships. The formulation below defines a smooth function of age and a linear regression term for PSQI_Comp5_Problems whose slope depends on age. Because the PSQI variable only takes on a discrete set of four values, we do not define a full smooth term for this one.

```
mod4 <- gamm4(Average_Thickness ~ s(Age) + s(Age, by = PSQI_Comp5_Problems, k = 6) + Sex + Site_Name,
              data = dat4, random = ~ (1|ID))
```

We set up a grid over which to plot the fits, and then compute and plot.

```
grid <- crossing(
  Site_Name = "Avanto",
  Sex = "Female",
  BL_Age = seq(from = 20, to = 90, by = 10),
  Interval = seq(from = 0, to = 5, by = 1),
  PSQI_Comp5_Problems = 0:3
) %>%
  mutate(Age = BL_Age + Interval)

prediction <- predict(mod4$gam, newdata = grid, se.fit = TRUE)

preds <- grid %>%
  mutate(
    fit = as.numeric(prediction$fit),
    se.fit = as.numeric(prediction$se.fit),
    BL_Age = factor(BL_Age),
    PSQI_Comp5_Problems = factor(PSQI_Comp5_Problems)
  )

p1 <- ggplot(preds, aes(x = Interval, y = fit,
                        ymin = fit + qnorm(.025) * se.fit,
                        ymax = fit + qnorm(.975) * se.fit,
                        color = PSQI_Comp5_Problems,
                        fill = PSQI_Comp5_Problems
                        )) +
  geom_line() +
  geom_ribbon(alpha = .2, color = NA) +
  facet_wrap(vars(BL_Age), scales = "free_y",
             labeller = as_labeller(function(x) paste("BL_Age =", x))) +
  theme_classic() +
  theme(legend.position = "bottom")

p1
```

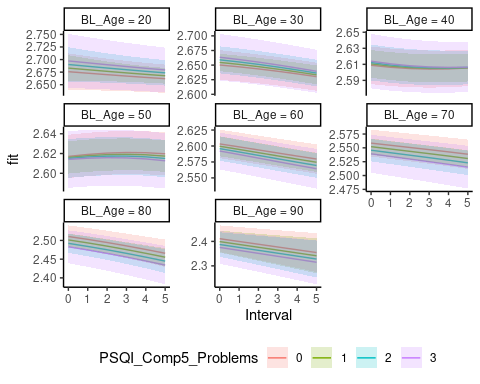

```
p2 <- ggplot(preds, aes(x = Interval, y = fit,
                        group = PSQI_Comp5_Problems,
                        color = PSQI_Comp5_Problems)) +
  geom_line() +
  facet_wrap(vars(BL_Age),
             labeller = as_labeller(function(x) paste("BL_Age =", x))) +
  theme_classic() +
  theme(legend.position = "bottom")

p2
```

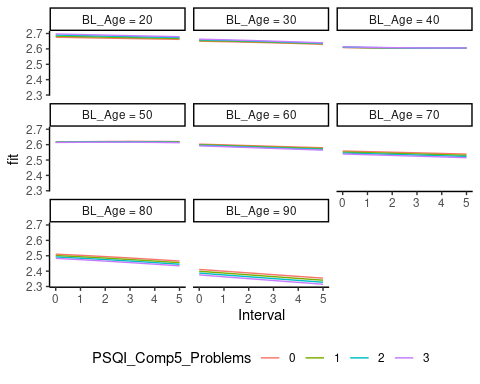

```
## Saving 5 x 4 in image
## Saving 5 x 4 in image
```

#### Controling for BMI

We next run to new models controling for BMI. Because the missingness of BMI might depend on other person characteristics, and definitely depends on cohort, we start by fitting the GAMM without controling for BMI on the dataset containing subjects with BMI values.

```
mod4_bmi_subjects <- gamm4(Average_Thickness ~ s(Age, k = 5) + s(Age, by = PSQI_Comp5_Problems, k = 5) +
                             Sex + Site_Name, data = dat4_bmi, random = ~(1|ID))
```

Next, we control for BMI by adding this as a linear predictor.

```
mod4_bmi_subjects_controled <- gamm4(Average_Thickness ~ s(Age, k = 5) + s(Age, by = PSQI_Comp5_Problems, k = 5) +
                             Sex + Site_Name + BMI, data = dat4_bmi, random = ~(1|ID))

summary(mod4_bmi_subjects_controled$gam)
##
## Family: gaussian
## Link function: identity
##
## Formula:
## Average_Thickness ~ s(Age, k = 5) + s(Age, by = PSQI_Comp5_Problems,
##     k = 5) + Sex + Site_Name + BMI
##
## Parametric coefficients:
##                   Estimate Std. Error t value Pr(>|t|)
## (Intercept)      2.5456553  0.0201549 126.304  < 2e-16 ***
## SexMale         -0.0251927  0.0060391  -4.172 3.12e-05 ***
## Site_NameBetula -0.0480049  0.0100170  -4.792 1.74e-06 ***
## Site_NameCamCAN  0.0599229  0.0083319   7.192 8.34e-13 ***
## Site_NamePrisma  0.0405750  0.0147684   2.747  0.00605 **
## Site_NameSkyra  -0.0405288  0.0067719  -5.985 2.47e-09 ***
## BMI              0.0016520  0.0007064   2.339  0.01943 *
## ---
## Signif. codes:  0 '***' 0.001 '**' 0.01 '*' 0.05 '.' 0.1 ' ' 1
##
## Approximate significance of smooth terms:
##                              edf Ref.df      F  p-value
## s(Age)                     3.334  3.334 18.113 3.31e-12 ***
## s(Age):PSQI_Comp5_Problems 2.000  2.000  0.428    0.652
## ---
## Signif. codes:  0 '***' 0.001 '**' 0.01 '*' 0.05 '.' 0.1 ' ' 1
##
## R-sq.(adj) =  0.243
## lmer.REML = -3550.5  Scale est. = 0.0072984  n = 2578
```

From the model output, the effects look very similar. We check this by plotting, and see that the fits are indistinguishable with and without controling for BMI.

```
grid <- crossing(
  Site_Name = "Avanto",
  Sex = "Female",
  BL_Age = seq(from = 20, to = 90, by = 10),
  Interval = seq(from = 0, to = 5, by = 1),
  PSQI_Comp5_Problems = 0:3, BMI = 20
) %>%
  mutate(Age = BL_Age + Interval)

prediction_bmi_subjects <- predict(mod4_bmi_subjects$gam, newdata = grid,
                                   type = "terms",
                                   terms = c("s(Age)", "s(Age):PSQI_Comp5_Problems")) %>%
  rowSums()
prediction_bmi_subjects_controled <- predict(mod4_bmi_subjects_controled$gam, newdata = grid,
                                   type = "terms",
                                   terms = c("s(Age)", "s(Age):PSQI_Comp5_Problems")) %>%
  rowSums()

preds <- grid %>%
  mutate(
    `without BMI` = prediction_bmi_subjects,
    `with BMI` = prediction_bmi_subjects_controled,
    BL_Age = factor(BL_Age),
    PSQI_Comp5_Problems = factor(PSQI_Comp5_Problems)
  ) %>%
  pivot_longer(cols = c("without BMI", "with BMI"))

ggplot(preds, aes(x = Interval, y = value,
                  group = paste(PSQI_Comp5_Problems, name),
                  color = PSQI_Comp5_Problems,
                  linetype = name)) +
  geom_line() +
  facet_wrap(vars(BL_Age), scales = "free_y",
             labeller = as_labeller(function(x) paste("BL_Age =", x))) +
  theme_classic() +
  theme(legend.position = "bottom") +
  labs(linetype = NULL)
```

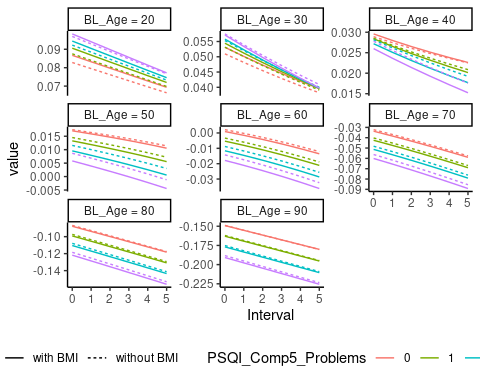

#### Controling for Depression

We next run to new models controling for Depression. Because the missingness of Depression might depend on other person characteristics, and definitely depends on cohort, we start by fitting the GAMM without controling for Depression on the dataset containing subjects with Depression values.

```
mod4_depression_subjects <- gamm4(Average_Thickness ~ s(Age, k = 5) + s(Age, by = PSQI_Comp5_Problems, k = 5) +
                             Sex + Site_Name, data = dat4_depression, random = ~(1|ID))
```

Next, we control for Depression by adding this as a linear predictor.

```
mod4_depression_subjects_controled <- gamm4(Average_Thickness ~ s(Age, k = 5) + s(Age, by = PSQI_Comp5_Problems, k = 5) +
                             Sex + Site_Name + Depression, data = dat4_depression, random = ~(1|ID))
```

From the model output, the effects look very similar. We check this by plotting, and see that the fits are indistinguishable with and without controling for Depression.

```
grid <- crossing(
  Site_Name = "Avanto",
  Sex = "Female",
  BL_Age = seq(from = 20, to = 90, by = 10),
  Interval = seq(from = 0, to = 5, by = 1),
  PSQI_Comp5_Problems = 0:3, Depression = 0
) %>%
  mutate(Age = BL_Age + Interval)

prediction_depression_subjects <- predict(mod4_depression_subjects$gam, newdata = grid,
                                   type = "terms",
                                   terms = c("s(Age)", "s(Age):PSQI_Comp5_Problems")) %>%
  rowSums()
prediction_depression_subjects_controled <- predict(mod4_depression_subjects_controled$gam, newdata = grid,
                                   type = "terms",
                                   terms = c("s(Age)", "s(Age):PSQI_Comp5_Problems")) %>%
  rowSums()

preds <- grid %>%
  mutate(
    `without Depression` = prediction_depression_subjects,
    `with Depression` = prediction_depression_subjects_controled,
    BL_Age = factor(BL_Age),
    PSQI_Comp5_Problems = factor(PSQI_Comp5_Problems)
  ) %>%
  pivot_longer(cols = c("without Depression", "with Depression"))

ggplot(preds, aes(x = Interval, y = value,
                  group = paste(PSQI_Comp5_Problems, name),
                  color = PSQI_Comp5_Problems,
                  linetype = name)) +
  geom_line() +
  facet_wrap(vars(BL_Age), scales = "free_y",
             labeller = as_labeller(function(x) paste("BL_Age =", x))) +
  theme_classic() +
  theme(legend.position = "bottom") +
  labs(linetype = NULL)
```

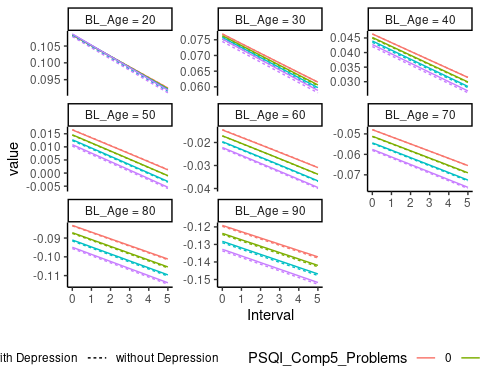

```
rm(dat4_bmi, dat4_depression, mod4_bmi_subjects,
   mod4_bmi_subjects_controled, mod4_depression_subjects,
   mod4_depression_subjects_controled
```

**5. Details about the memory analyses**

Memory and Sleep

### Sample Descriptives

We have the following memory variables, scaled to zero mean and unit standard deviation.

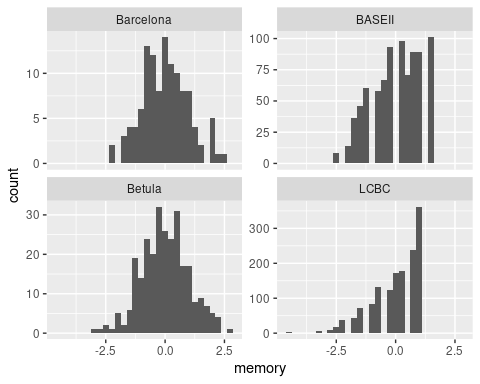

We have the following sample characteristics.

| Study | Unique subjects | Total observations | Mean baseline age | Age range |
| --- | --- | --- | --- | --- |
| Barcelona | 39 | 116 | 69 | 64 - 81 |
| BASEII | 512 | 830 | 62 | 14 - 92 |
| Betula | 138 | 276 | 65 | 55 - 85 |
| LCBC | 730 | 1480 | 49 | 19 - 89 |

The plot below shows the actual data for memory versus age. Note the ceiling effect in the LCBC data, which uses CVLT 30 minutes free recall and in the BASEII data.

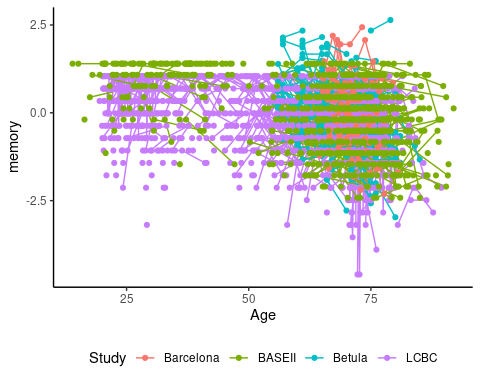

We now fit models relating PSQI to memory change. We use the following components, separately. Medication (Comp 6) is not included.

```
PSQI_comps <- c("PSQI_Comp1_Quality", "PSQI_Comp2_Latency",
                "PSQI_Comp3_Duration", "PSQI_Comp4_Efficiency",
                "PSQI_Comp5_Problems", "PSQI_Comp7_Tired",
                "PSQI_Global")
```

### Effect of Sleep on Memory Change

Models without age interaction. We use a nested random effects structure with subjects (IDs) within studies. In addition, we allow the effect of time and sleep to vary between studies, by including these as random effects per study. Given the nonlinear nature of the age-memory relation, we use a smooth term for baseline age.

```
library(gamm4)
models <- map(PSQI_comps, function(component){
  gamm4(memory ~ s(BL_Age, bs = "cr") + Time * sleep + Sex,
        data = filter(memory_data, component == !!component),
        random = ~(Time * sleep|Study) + (1|Study:ID))
})
names(models) <- PSQI_comps
```

Below is a plot of the estimated effect of baseline age on memory. It is almost identical for all models.

plot(models[[1]]$gam, select = 1, scale = 0)

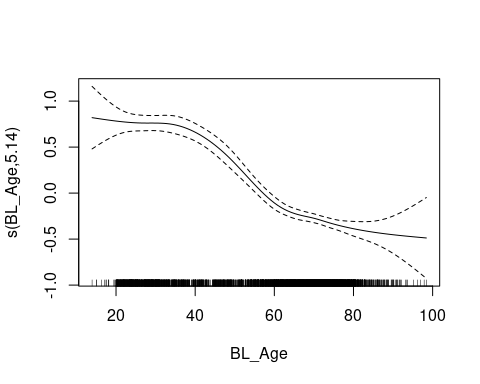

```
vars <- c("Time", "sleep", "Time:sleep")
df <- map_dfr(models, function(mod){
  tab <- summary(mod$gam)$p.table

  tibble(
    var = !!vars,
    estimates = tab[vars, "Estimate"],
    estimate_se = tab[vars, "Std. Error"],
    pval = tab[vars, "Pr(>|t|)"]
  )
}, .id = "Component")
df_ran <- map_dfr(models, function(mod){
  tab <- ranef(mod$mer)$Study[, vars]
  tab$Study <- rownames(tab)
  fixef(mod$mer)

  as_tibble(tab) %>%
    pivot_longer(cols = -Study, values_to = "ranef", names_to = "var") %>%
    mutate(
      fixef = fixef(mod$mer)[paste0("X", var)],
      estimates = fixef + ranef
    )
}, .id = "Component")
```

Below are plots of the model coefficients. The error bars show 95 % confidence intervals.

```
ggplot(df, aes(x = Component, y = estimates)) +
  geom_point() +
  geom_errorbar(width = .3, aes(ymin = estimates + qnorm(.025) * estimate_se,
               ymax = estimates + qnorm(.975) * estimate_se)) +
  geom_hline(yintercept = 0, linetype = "dashed") +
  facet_wrap(vars(var), scales = "free_y") +
  theme_classic() +
  theme(axis.text.x = element_text(angle = 90))
```

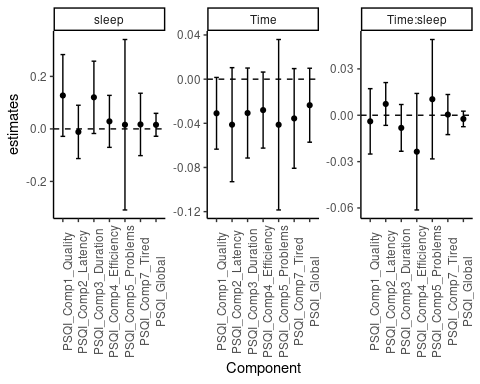

In the plot below, the random effect per study has been added to each plot, showing the heterogeneity.

```
ggplot(df, aes(x = Component, y = estimates)) +
  geom_point() +
  geom_point(data = df_ran, aes(color = Study)) +
  geom_errorbar(width = .3, aes(ymin = estimates + qnorm(.025) * estimate_se,
               ymax = estimates + qnorm(.975) * estimate_se)) +
  geom_hline(yintercept = 0, linetype = "dashed") +
  facet_wrap(vars(var), scales = "free_y") +
  theme_classic() +
  theme(axis.text.x = element_text(angle = 90))
```

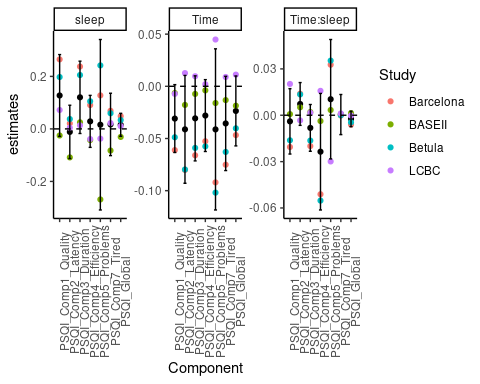

The table below shows the p-values for each of the model coefficients.

```
df %>%
  select(-estimates, -estimate_se) %>%
  pivot_wider(names_from = var, values_from = pval) %>%
  knitr::kable(digits = 3)
```

| Component | Time | sleep | Time:sleep |
| --- | --- | --- | --- |
| PSQI_Comp1_Quality | 0.062 | 0.109 | 0.715 |
| PSQI_Comp2_Latency | 0.118 | 0.826 | 0.301 |
| PSQI_Comp3_Duration | 0.140 | 0.087 | 0.290 |
| PSQI_Comp4_Efficiency | 0.111 | 0.569 | 0.220 |
| PSQI_Comp5_Problems | 0.295 | 0.923 | 0.598 |
| PSQI_Comp7_Tired | 0.122 | 0.777 | 0.945 |
| PSQI_Global | 0.167 | 0.482 | 0.354 |

### Effect of Sleep and Age on Memory Change

The models are as before, except that we now also include the three-way interaction between time, sleep, and baseline age. Note that Time * sleep * BL_Age - BL_Age specifies that we want the three-way interaction and all lower-order terms, except BL_Age, since the latter is taken care of by the smooth term.

```
library(gamm4)
models <- map(PSQI_comps, function(component){
  gamm4(memory ~ s(BL_Age, bs = "cr") + Time * sleep * BL_Age - BL_Age + Sex,
        data = filter(memory_data, component == !!component),
        random = ~(Time * sleep * BL_Age|Study) + (1|Study:ID))
})
names(models) <- PSQI_comps
vars <- c("Time", "sleep", "Time:sleep", "Time:BL_Age", "sleep:BL_Age",
          "Time:sleep:BL_Age")
df <- map_dfr(models, function(mod){
  tab <- summary(mod$gam)$p.table

  tibble(
    var = !!vars,
    estimates = tab[vars, "Estimate"],
    estimate_se = tab[vars, "Std. Error"],
    pval = tab[vars, "Pr(>|t|)"]
  )
}, .id = "Component")
df_ran <- map_dfr(models, function(mod){
  tab <- ranef(mod$mer)$Study[, vars]
  tab$Study <- rownames(tab)
  fixef(mod$mer)

  as_tibble(tab) %>%
    pivot_longer(cols = -Study, values_to = "ranef", names_to = "var") %>%
    mutate(
      fixef = fixef(mod$mer)[paste0("X", var)],
      estimates = fixef + ranef
    )
}, .id = "Component")
```

Below are plots of the model coefficients. The error bars show 95 % confidence intervals.

```
ggplot(df, aes(x = Component, y = estimates)) +
  geom_point() +
  geom_errorbar(width = .3, aes(ymin = estimates + qnorm(.025) * estimate_se,
               ymax = estimates + qnorm(.975) * estimate_se)) +
  geom_hline(yintercept = 0, linetype = "dashed") +
  facet_wrap(vars(var), scales = "free_y") +
  theme_classic() +
  theme(axis.text.x = element_text(angle = 90))
```

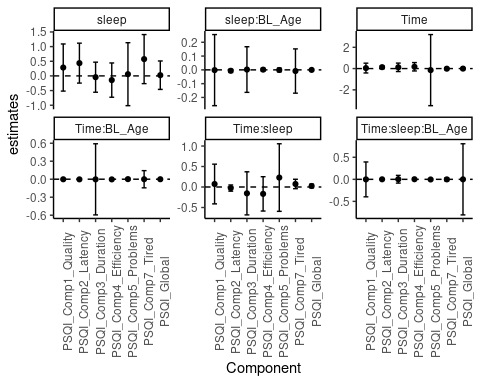

We plot again without confidence intervals, for visibility.

```
ggplot(df, aes(x = Component, y = estimates)) +
  geom_point() +
  geom_hline(yintercept = 0, linetype = "dashed") +
  facet_wrap(vars(var), scales = "free_y") +
  theme_classic() +
  theme(axis.text.x = element_text(angle = 90))
```

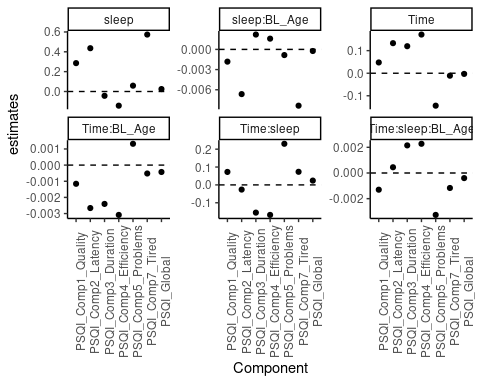

In the plot below, the random effect per study has been added to each plot, showing the heterogeneity.

```
ggplot(df, aes(x = Component, y = estimates)) +
  geom_point() +
  geom_point(data = df_ran, aes(color = Study)) +
  geom_hline(yintercept = 0, linetype = "dashed") +
  facet_wrap(vars(var), scales = "free_y") +
  theme_classic() +
  theme(axis.text.x = element_text(angle = 90))
```

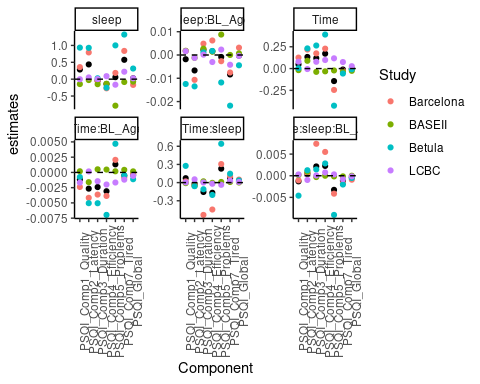

The table below shows the p-values for each of the model coefficients.

```
df %>%
  select(-estimates, -estimate_se) %>%
  pivot_wider(names_from = var, values_from = pval) %>%
  knitr::kable(digits = 3)
```

| Component | Time | sleep | Time:sleep | Time:BL_Age | sleep:BL_Age | Time:sleep:BL_Age |
| --- | --- | --- | --- | --- | --- | --- |
| PSQI_Comp1_Quality | 0.839 | 0.486 | 0.769 | 0.794 | 0.989 | 0.995 |
| PSQI_Comp2_Latency | 0.049 | 0.209 | 0.500 | 0.091 | 0.269 | 0.723 |
| PSQI_Comp3_Duration | 0.554 | 0.868 | 0.564 | 0.994 | 0.979 | 0.962 |
| PSQI_Comp4_Efficiency | 0.394 | 0.632 | 0.433 | 0.389 | 0.762 | 0.401 |
| PSQI_Comp5_Problems | 0.933 | 0.915 | 0.584 | 0.839 | 0.903 | 0.576 |
| PSQI_Comp7_Tired | 0.829 | 0.179 | 0.212 | 0.994 | 0.919 | 0.948 |
| PSQI_Global | 0.945 | 0.921 | 0.280 | 0.697 | 0.951 | 0.999 |

**6. Quantification of Aβ**

**Supplemental Information on quantification of *Aβ***

*LCBC Aβ sample PET*

Participants underwent an ^18^F-flutemetamol-PET scan, sensitive to Aβ accumulation^46^. Images were acquired on a General Electric Discovery PET/CT 690 scanner at Aleris Hospital and Radiology, Oslo, Norway. A low-dose computerized tomography scan was first performed for subsequent attenuation correction of the PET scan. Participants were injected with 200±20 MBq ^18^F-flutemetamol as a bolus, and examined 90 minutes after the injection. Three-dimensional dynamic data was acquired in list mode for 20 minutes, with the following parameters: 47 image planes, voxel size = 1.33 mm x 1.33 mm x 3.27 mm, field of view = 256 mm. The images were reconstructed using the VUEPoint HD Sharp iterative reconstruction algorithm (including time-of-flight). This algorithm adds resolution recovery in an iterative reconstruction loop by incorporating information about the PET detector response, which improves resolution and contrast recovery compared with traditional analytic methods^47^. We used 4 iterations, 16 subsets, and a full width at half maximum of the Gaussian post-filter of 3 mm. As we were interested in the gross tracer uptake, we binned the data into a single frame, and submitted this static PET image to further pre-processing and value extraction. For PET pre-processing, we used PetSurfer, a set of tools within the FreeSurfer suite, for partial volume correction. Specifically, for each participant, we registered the static PET image to the anatomical T1-weighted image using boundary-based registration^48^. This registration was inverted to get a high-resolution segmentation (upsample factor = 2) from the high-resolution MRI space in PET space, and simultaneously perform the partial volume correction with the region-based voxel-wise method, as recommended when using surface-based vertex-wise analyses^49,50^. This procedure yielded PET signal for each vertex on the cortical surface. The PET signal in each vertex was divided by the mean signal of the cerebellum cortex to obtain standardized uptake value ratios (SUVR)^51^, and subjected to general linear model analyses. The initial vertex-wise analysis showed no significant association surviving proper correction for multiple comparisons (se figure below for unthresholded statistical maps). Thus, we parcellated the cortex into 34 regions in each hemisphere according to the Desikan-Killiany scheme, and ran a principal component analysis to extract a major amyloid factor. This factor explained 66.7% of the variance, and was entered into subsequent statistical analyses.

| 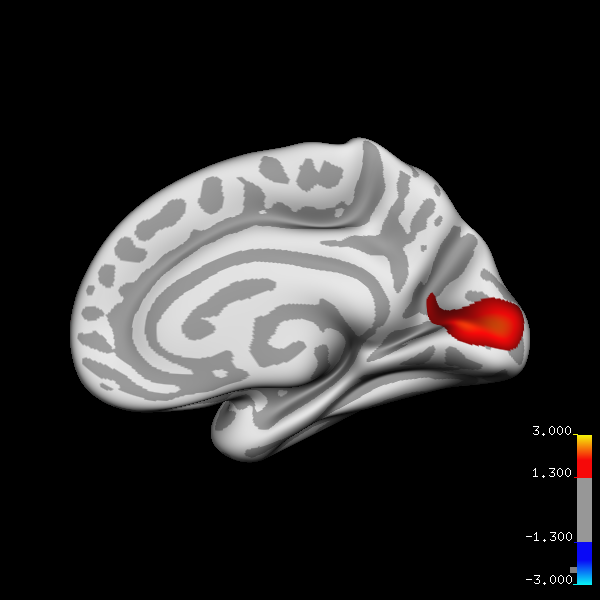 | 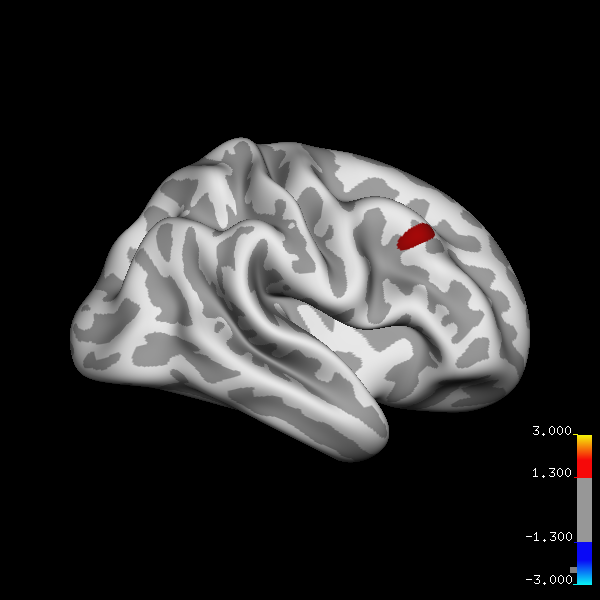 |
| --- | --- |

*Supplemental Information of LCBC PET data reductions*

The Aβ PET values were averaged in each of the 34 cortical regions in each hemisphere in the Desikan-Killiany atlas. The values were entered into a principal component analysis to obtain one main PET component. The results of this analysis is provided below.

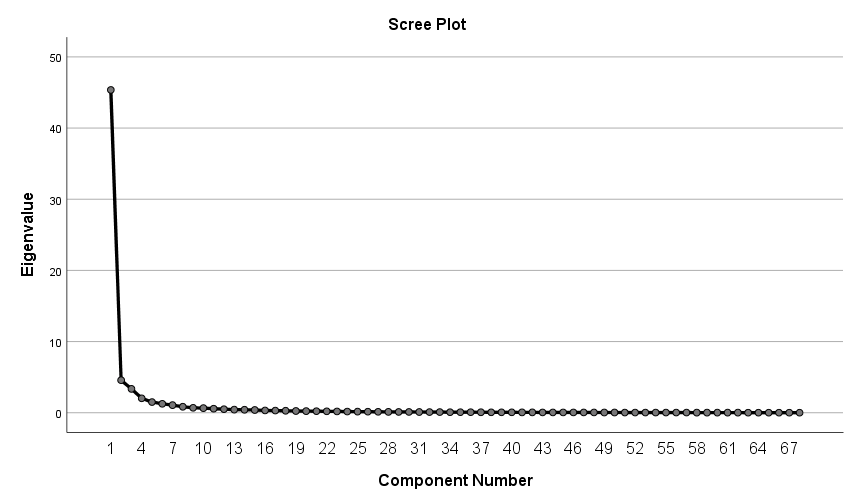

| **Total Variance Explained** | | | |
| --- | --- | --- | --- |
| Component | Extraction Sums of Squared Loadings | | |
|  | Total | % of Variance | Cumulative % |
| 1 | 45,359 | 66,704 | 66,704 |
| Extraction Method: Principal Component Analysis. | | | |

| **Component Matrix^a^** | |
| --- | --- |
|  | Component |
|  | 1 |
| PET_aparc_lh_supramargil | ,936 |
| PET_aparc_lh_insula | ,933 |
| PET_aparc_rh_precuneus | ,932 |
| PET_aparc_rh_inferiorparietal | ,931 |
| PET_aparc_lh_lateralorbitofrontal | ,929 |
| PET_aparc_rh_posteriorcingulate | ,928 |
| PET_aparc_lh_rostralmiddlefrontal | ,928 |
| PET_aparc_lh_posteriorcingulate | ,927 |
| PET_aparc_rh_parsopercularis | ,926 |
| PET_aparc_rh_insula | ,924 |
| PET_aparc_rh_superiorfrontal | ,923 |
| PET_aparc_rh_rostralmiddlefrontal | ,922 |
| PET_aparc_lh_precuneus | ,921 |
| PET_aparc_lh_isthmuscingulate | ,920 |
| PET_aparc_lh_superiorfrontal | ,918 |
| PET_aparc_lh_inferiorparietal | ,912 |
| PET_aparc_lh_inferiortemporal | ,911 |
| PET_aparc_lh_parsopercularis | ,910 |
| PET_aparc_rh_lateralorbitofrontal | ,909 |
| PET_aparc_rh_medialorbitofrontal | ,908 |
| PET_aparc_lh_rostralanteriorcingulate | ,907 |
| PET_aparc_rh_parsorbitalis | ,902 |
| PET_aparc_lh_medialorbitofrontal | ,902 |
| PET_aparc_lh_superiortemporal | ,900 |
| PET_aparc_rh_isthmuscingulate | ,895 |
| PET_aparc_rh_rostralanteriorcingulate | ,893 |
| PET_aparc_rh_supramargil | ,892 |
| PET_aparc_rh_inferiortemporal | ,892 |
| PET_aparc_lh_parsorbitalis | ,891 |
| PET_aparc_lh_middletemporal | ,891 |
| PET_aparc_lh_bankssts | ,890 |
| PET_aparc_rh_superiortemporal | ,886 |
| PET_aparc_rh_bankssts | ,885 |
| PET_aparc_rh_parstriangularis | ,882 |
| PET_aparc_lh_parstriangularis | ,877 |
| PET_aparc_rh_caudalmiddlefrontal | ,876 |
| PET_aparc_rh_caudalanteriorcingulate | ,876 |
| PET_aparc_lh_caudalmiddlefrontal | ,868 |
| PET_aparc_rh_middletemporal | ,863 |
| PET_aparc_lh_fusiform | ,847 |
| PET_aparc_lh_caudalanteriorcingulate | ,847 |
| PET_aparc_rh_superiorparietal | ,835 |
| PET_aparc_lh_parahippocampal | ,835 |
| PET_aparc_rh_fusiform | ,834 |
| PET_aparc_lh_superiorparietal | ,833 |
| PET_aparc_lh_frontalpole | ,825 |
| PET_aparc_rh_frontalpole | ,810 |
| PET_aparc_lh_postcentral | ,761 |
| PET_aparc_rh_parahippocampal | ,741 |
| PET_aparc_rh_postcentral | ,725 |
| PET_aparc_lh_paracentral | ,722 |
| PET_aparc_lh_transversetemporal | ,706 |
| PET_aparc_rh_paracentral | ,692 |
| PET_aparc_lh_precentral | ,669 |
| PET_aparc_rh_precentral | ,663 |
| PET_aparc_rh_lateraloccipital | ,662 |
| PET_aparc_lh_temporalpole | ,662 |
| PET_aparc_lh_lateraloccipital | ,643 |
| PET_aparc_rh_transversetemporal | ,638 |
| PET_aparc_rh_temporalpole | ,635 |
| PET_aparc_lh_pericalcarine | ,498 |
| PET_aparc_rh_pericalcarine | ,497 |
| PET_aparc_lh_entorhil | ,477 |
| PET_aparc_rh_lingual | ,466 |
| PET_aparc_rh_entorhil | ,458 |
| PET_aparc_lh_lingual | ,456 |
| PET_aparc_lh_cuneus | ,405 |
| PET_aparc_rh_cuneus | ,390 |
| Extraction Method: Principal Component Analysis. | |
| a. 1 components extracted. | |

*LCBC Aβ sample CSF*

CSF was collected in polypropylene tubes, centrifuged, aliquoted and stored at – 80° C, as described elsewhere ^45^. Samples were sent on dry ice to the laboratory, and CSF Aβ42 was determined using INNOTEST enzyme-linked immunosorbent assays (ELISA; Fujirebio, Ghent, Belgium) at Sahlgrenska University Hospital (Mölndal, Sweden) by board-certified laboratory technicians, as previously described ^45^.

*ADNI*

Participants from ADNI were chosen based on the following algorithm. All participant with available data on sleep and Aβ were chosen for analysis. Since both CSF Aβ42and ^18^F-florbetapir PET were available for multiple participants, the following algorithm was used: 1) Assign participants with only one Aβ measure to their corresponding group, 2) For participants with more Aβ measures, assign the participant to the dataset with more observations, 3) For the remaining participants, randomly assign participants to the CSF or the PET group. This resulted in one group of 846 (1937 obervations) participants with Aβ from PET and one group of 870 participants (1717 observations) with Aβ from CSF. Quantification of Aβ from PET and CSF are described at the ADNI page (adni.loni.usc.edu), specifically <http://adni.loni.usc.edu/methods/pet-analysis-method/pet-analysis/> (PET analysis), <http://adni.loni.usc.edu/methods/pet-analysis-method/pet-analysis/#pet-pre-processing-container> (Pet preprocessing) and <http://adni.loni.usc.edu/methods/> and (CSF analysis). In short, for PET the following protocol was run: Florbetapir (Amyvid): 370 MBq (10.0 mCi) ± 10%, 20 min (4X5min frames) acquisition at 50-70 min post-injection.

**7. Virtual histology analyses**

In stage 1, we calculated the mean Spearman correlation between each of the six donor’s profiles and the median (group) profile for each gene. Using this metric, the median profile provides a good approximation across the donors for 39.7% of the assayed genes (rho > 0.446 corresponding to one-sided p < 0.05 derived from random simulations of donor expression profiles) (French and Paus, 2015). In stage 2, we relied on the BrainSpan Atlas, which provides gene expression data in the developing human brain ([www.brainspan.org](http://www.brainspan.org)) (Miller et al., 2014) as previously implemented (Shin et al., 2018b). We limited the samples to age > 12 years (n = 9 donors) and downloaded gene expression values obtained in 11 cortical regions included in the BrainSpan atlas that are homologous to those on the Desikan-Killiany parcellation employed in the Allen Human Brain Atlas. Next, we compared the similarity between the regional profile between the Allen Human Brain Atlas and the BrainSpan Atlas across the 11 cortical regions available in both atlases. Only genes that showed a correlation between the two profiles higher than r = 0.52 (one-sided test P < 0.05) were retained. Finally we converted mouse genes to human gene symbols as implemented in the *“homologene”* R package (O’Leary et al., 2016).

**8. Complete listing of ADNI researchers**

A complete listing of Alzheimer’s Disease Neuroimaging Initiative (ADNI) investigators is as follows: ADNI I, GO, II, and III Leadership and Infrastructure: Principal Investigator: Michael W. Weiner, MD, University of California, San Francisco; ATRI PI and Director of Coordinating Center Clinical Core: Paul Aisen, MD, University of Southern California; Executive Committee: Michael Weiner, MD, University of California, San Francisco; Paul Aisen, MD, University of Southern California; Ronald Petersen, MD, PhD, Mayo Clinic, Rochester; Clifford R. Jack, Jr, MD, Mayo Clinic, Rochester; William Jagust, MD, University of California, Berkeley; John Q. Trojanowki, MD, PhD, University of Pennsylvania; Arthur W. Toga, PhD, University of Southern California; Laurel Beckett, PhD, University of California, Davis; Robert C. Green, MD, MPH, Brigham and Women’s Hospital/Harvard Medical School; Andrew J. Saykin, PsyD, Indiana University; John Morris, MD, Washington University, St Louis; Leslie M. Shaw, University of Pennsylvania; ADNI External Advisory Board: Zaven Khachaturian, PhD, Prevent Alzheimer’s Disease 2020 (chair); Greg Sorensen, MD, Siemens; Maria Carrillo, PhD, Alzheimer’s Association; Lew Kuller, MD, University of Pittsburgh; Marc Raichle, MD, Washington University, St Louis; Steven Paul, MD, Cornell University; Peter Davies, MD, Albert Einstein College of Medicine of Yeshiva University; Howard Fillit, MD, AD Drug Discovery Foundation; Franz Hefti, PhD, Acumen Pharmaceuticals; David Holtzman, MD, Washington University, St Louis; M. Marcel Mesulam, MD, Northwestern University; William Potter, MD, National Institute of Mental Health; Peter Snyder, PhD, Brown University; ADNI 3 Private Partner Scientific Board: Veronika Logovinsky, MD, PhD, Eli Lilly (chair); Data and Publications Committee: Robert C. Green, MD, MPH, Brigham and Women’s Hospital/Harvard Medical School; (chair); Resource Allocation Review Committee: Tom Montine, MD, PhD, University of Washington (chair); Clinical Core Leaders: Ronald Petersen, MD, PhD, Mayo Clinic, Rochester (core principal investigator); Paul Aisen, MD, University of Southern California; Clinical Informatics and Operations: Gustavo Jimenez, MBS, University of Southern California; Michael Donohue, PhD, University of Southern California; Devon Gessert, BS, University of Southern California; Kelly Harless, BA, University of Southern California; Jennifer Salazar, MBS, University of Southern California; Yuliana Cabrera, BS, University of Southern California; Sarah Walter, MSc, University of Southern California; Lindsey Hergesheimer, BS, University of Southern California; Biostatistics Core Leaders and Key Personnel: Laurel Beckett, PhD, University of California, Davis (core principal investigator); Danielle Harvey, PhD, University of California, Davis; Michael Donohue, PhD, University of California, San Diego; MRI Core Leaders and Key Personnel: Clifford R. Jack, Jr, MD, Mayo Clinic, Rochester (core principal investigator); Matthew Bernstein, PhD, Mayo Clinic, Rochester; Nick Fox, MD, University of London; Paul Thompson, PhD, UCLA School of Medicine; Norbert Schuff, PhD, University of California, San Francisco, MRI; Charles DeCarli, MD, University of California, Davis; Bret Borowski, RT, Mayo Clinic; Jeff Gunter, PhD, Mayo Clinic; Matt Senjem, MS, Mayo Clinic; Prashanthi Vemuri, PhD, Mayo Clinic; David Jones, MD, Mayo Clinic; Kejal Kantarci, Mayo Clinic; Chad Ward Mayo Clinic; PET Core Leaders and Key Personnel: William Jagust, MD, University of California, Berkeley (core principal investigator); Robert A. Koeppe, PhD, University of Michigan; Norm Foster, MD, University of Utah; Eric M. Reiman, MD, Banner Alzheimer’s Institute; Kewei Chen, PhD, Banner Alzheimer’s Institute; Chet Mathis, MD, University of Pittsburgh; Susan Landau, PhD, University of California, Berkeley; Neuropathology Core Leaders: John C. Morris, MD, Washington University, St Louis; Nigel J. Cairns, PhD, FRCPath, Washington University, St Louis; Erin Franklin, MS, CCRP Washington University, St Louis; Lisa Taylor-Reinwald, BA, HTL, Washington University, St Louis (past investigator); Biomarkers Core Leaders and Key Personnel: Leslie M. Shaw, PhD, University of Pennsylvania School of Medicine; John Q. Trojanowki, MD, PhD, University of Pennsylvania School of Medicine; Virginia Lee, PhD, MBA, University of Pennsylvania School of Medicine; Magdalena Korecka, PhD, University of Pennsylvania School of Medicine; Michal Figurski, PhD, University of Pennsylvania School of Medicine; Informatics Core Leaders and Key Personnel: Arthur W. Toga, PhD, University of Southern California (core principal investigator); Karen Crawford, University of Southern California; Scott Neu, PhD, University of Southern California; Genetics Core Leaders and Key Personnel: Andrew J. Saykin, PsyD, Indiana University; Tatiana M. Foroud, PhD, Indiana University; Steven Potkin, MD, University of California, Irvine; Li Shen, PhD, Indiana University; Kelley Faber, MS, CCRC, Indiana University; Sungeun Kim, PhD, Indiana University; Kwangsik Nho, PhD, Indiana University; Initial Concept Planning & Development: Michael W. Weiner, MD, University of California, San Francisco; Lean Thal, MD, University of California, San Diego; Zaven Khachaturian, PhD, Prevent Alzheimer’s Disease 2020; Early Project Proposal Development: Leon Thal, MD, University of California, San Diego; Neil Buckholtz, National Institute on Aging; Michael W. Weiner, MD, University of California, San Francisco; Peter J. Snyder, PhD, Brown University; William Potter, MD, National Institute of Mental Health; Steven Paul, MD, Cornell University; Marilyn Albert, PhD, Johns Hopkins University; Richard Frank, MD, PhD, Richard Frank Consulting; Zaven Khachaturian, PhD, Prevent Alzheimer’s Disease 2020; National Institute on Aging: John Hsiao, MD, National Institute on Aging; Investigators by Site Oregon Health & Science University: Joseph Quinn, MD; Lisa C. Silbert, MD; Betty Lind, BS; Jeffrey A. Kaye, MD (past investigator); Raina Carter, BA (past investigator); Sara Dolen, BS (past investigator); University of Southern California: Lon S. Schneider, MD; Sonia Pawluczyk, MD; Mauricio Becerra, BS; Liberty Teodoro, RN; Bryan M. Spann, DO, PhD (past investigator); University of California, San Diego: James Brewer, MD, PhD; Helen Vanderswag, RN; Adam Fleisher, MD (past investigator); University of Michigan: Jaimie Ziolkowski, MA, BS, TLL; Judith L. Heidebrink, MD, M; Joanne L. Lord, LPN, BA, CCRC (past investigator); Mayo Clinic, Rochester: Ronald Petersen, MD, PhD; Sara S. Mason, RN; Colleen S. Albers, RN; David Knopman, MD; Kris Johnson, RN (past investigator); Baylor College of Medicine: Javier Villanueva-Meyer, MD; Valory Pavlik, PhD; Nathaniel Pacini, MA; Ashley Lamb, MA; Joseph S. Kass, MD, LD, FAAN; Rachelle S. Doody, MD, PhD (past investigator); Victoria Shibley, MS (past investigator); Munir Chowdhury, MBBS, MS (past investigator); Susan Rountree, MD (past investigator); Mimi Dang, MD (past investigator); Columbia University Medical Center: Yaakov Stern, PhD; Lawrence S. Honig, MD, PhD; Karen L. Bell, MD; Randy Yeh, MD. Washington University, St Louis: Beau Ances, MD, PhD, MSc; John C. Morris, MD; David Winkfield, BS; Maria Carroll, RN, MSN, GCNS-BC; Angela Oliver, RN, BSN, MSG; Mary L. Creech, RN, MSW (past investigator); Mark A. Mintun, MD (past investigator); Stacy Schneider, APRN, BC, GNP (past investigator); University of Alabama, Birmingham: Daniel Marson, JD, PhD; David Geldmacher, MD; Marissa Natelson Love, MD; Randall Griffith, PhD, ABPP (past investigator); David Clark, MD (past investigator); John Brockington, MD (past investigator); Mount Sinai School of Medicine: Hillel Grossman, MD; Effie Mitsis, PhD (past investigator); Rush University Medical Center: Raj C. Shah, MD; Melissa Lamar, PhD; Patricia Samuels; Wien Center: Ranjan Duara, MD; Maria T. Greig-Custo, MD; Rosemarie Rodriguez, PhD; Johns Hopkins University: Marilyn Albert, PhD; Chiadi Onyike, MD; Daniel D’Agostino II, BS; Stephanie Kielb, BS (past investigator); New York University: Martin Sadowski, MD, PhD; Mohammed O. Sheikh, MD; Jamika Singleton-Garvin, CCRP; Anaztasia Ulysse Mrunalini Gaikwad; Duke University Medical Center: P. Murali Doraiswamy, MBBS, FRCP; Jeffrey R. Petrella, MD; Olga James, MD; Salvador Borges-Neto, MD; Terence Z. Wong, MD (past investigator); Edward Coleman (past investigator); University of Pennsylvania: Jason H. Karlawish, MD; David A. Wolk, MD; Sanjeev Vaishnavi, MD; Christopher M. Clark, MD (past investigator); Steven E. Arnold, MD (past investigator); University of Kentucky: Charles D. Smith, MD; Greg Jicha, MD; Peter Hardy, PhD; Riham El Khouli, MD; Elizabeth Oates, MD; Gary Conrad, MD; University of Pittsburgh: Oscar L. Lopez, MD; MaryAnn Oakley, MA; Donna M. Simpson, CRNP, MPH; University of Rochester Medical Center: Anton P. Porsteinsson, MD; Kim Martin, RN; Nancy Kowalksi, MS, RNC; Melanie Keltz, RN; Bonnie S. Goldstein, MS, NP (past investigator); Kelly M. Makino, BS (past investigator); M. Saleem Ismail, MD (past investigator); Connie Brand, RN (past investigator); University of California Irvine IMIND: Gaby Thai, MD; Aimee Pierce, MD; Beatriz Yanez, RN; Elizabeth Sosa, PhD; Megan Witbracht, PhD; University of Texas Southwestern Medical School: Kyle Womack, MD; Dana Mathews, MD, PhD; Mary Quiceno, MD; Emory University: Allan I. Levey, MD, PhD; James J. Lah, MD, PhD; Janet S. Cellar, DNP, PMHCNS-BC University of Kansas, Medical Center: Jeffrey M. Burns, MD; Russell H. Swerdlow, MD; William M. Brooks, PhD; University of California, Los Angeles: Ellen Woo, PhD; Daniel H.S. Silverman, MD, PhD; Edmond Teng, MD, PhD; Sarah Kremen, MD; Liana Apostolova, MD (past investigator); Kathleen Tingus, PhD (past investigator); Po H. Lu, PsyD (past investigator); George Bartzokis, MD (past investigator); Mayo Clinic, Jacksonville: Neill R Graff-Radford, MBBCH, FRCP (London) Francine Parfitt, MSH, CCRC Kim Poki-Walker, BA; Indiana University: Martin R. Farlow, MD; Ann Marie Hake, MD; Brandy R. Matthews, MD (past investigator); Jared R. Brosch, MD; Scott Herring, RN, CCRC Yale University School of Medicine: Christopher H. van Dyck, MD; Richard E. Carson, PhD; Pradeep Varma, MD; McGill University, Montreal-Jewish General Hospital: Howard Chertkow, MD; Howard Bergman, MD; Chris Hosein, MEd, Sunnybrook Health Sciences, Ontario: Sandra Black, MD, FRCPC Bojana Stefanovic, PhD; Chris (Chinthaka) Heyn, BSC, PhD, MD, FRCPC U.B.C. Clinic for AD & Related Disorders: Ging-Yuek Robin Hsiung, MD, MHSc, FRCPC Benita Mudge, BS; Vesna Sossi, PhD; Howard Feldman, MD, FRCPC (past investigator); Michele Assaly, MA (past investigator); Cognitive Neurology - St. Joseph's, Ontario: Elizabeth Finger, MD; Stephen Pasternack, MD, PhD; William Pavlosky, MD; Irina Rachinsky, MD (past investigator); Dick Drost, PhD (past investigator); Andrew Kertesz, MD (past investigator); Cleveland Clinic Lou Ruvo Center for Brain Health: Charles Bernick, MD, MPH; Donna Munic, PhD; Northwestern University: Marek-Marsel Mesulam, MD; Emily Rogalski, PhD; Kristine Lipowski, MA; Sandra Weintraub, PhD; Borna Bonakdarpour, MD; Diana Kerwin, MD (past investigator); Chuang-Kuo Wu, MD, PhD (past investigator); Nancy Johnson, PhD (past investigator); Premiere Research Inst (Palm Beach Neurology): Carl Sadowsky, MD; Teresa Villena, MD; Georgetown University Medical Center: Raymond Scott Turner, MD, PhD; Kathleen Johnson, NP Brigid Reynolds, NP Brigham and Women's Hospital: Reisa A. Sperling, MD; Keith A. Johnson, MD; Gad A. Marshall, MD; Stanford University: Jerome Yesavage, MD; Joy L. Taylor, PhD; Steven Chao, MD, PhD; Barton Lane, MD (past investigator); Allyson Rosen, PhD (past investigator); Jared Tinklenberg, MD (past investigator); Banner Sun Health Research Institute: Edward Zamrini, MD; Christine M. Belden, PsyD; Sherye A. Sirrel, CCRC Boston University: Neil Kowall, MD; Ronald Killiany, PhD; Andrew E. Budson, MD; Alexander Norbash, MD (past investigator); Patricia Lynn Johnson, BA (past investigator); Howard University: Thomas O. Obisesan, MD, MPH; Ntekim E. Oyonumo, MD, PhD; Joanne Allard, PhD; Olu Ogunlana, BPharm; Case Western Reserve University: Alan Lerner, MD; Paula Ogrocki, PhD; Curtis Tatsuoka, PhD; Parianne Fatica, BA, CCRC; University of California, Davis – Sacramento: Evan Fletcher, PhD; Pauline Maillard, PhD; John Olichney, MD; Charles DeCarli, MD; Owen Carmichael, PhD (past investigator); Neurological Care of CNY: Smita Kittur, MD (past investigator); Parkwood Institute: Michael Borrie, MB ChB; T-Y Lee, PhD; Rob Bartha, PhD; University of Wisconsin: Sterling Johnson, PhD; Sanjay Asthana, MD; Cynthia M. Carlsson, MD, MS; Banner Alzheimer's Institute: Pierre Tariot, MD; Anna Burke, MD; Joel Hetelle, BS; Kathryn DeMarco, BS; Nadira Trncic, MD, PhD, CCRC (past investigator); Adam Fleisher, MD (past investigator); Stephanie Reeder, BA (past investigator); Dent Neurologic Institute: Vernice Bates, MD; Horacio Capote, MD; Michelle Rainka, PharmD, CCRP; Ohio State University: Douglas W. Scharre, MD; Maria Kataki, MD, PhD; Rawan Tarawneh, MD; Albany Medical College: Earl A. Zimmerman, MD; Dzintra Celmins, MD; David Hart, MD; Hartford Hospital, Olin Neuropsychiatry Research Center: Godfrey D. Pearlson, MD; Karen Blank, MD; Karen Anderson, RN; Dartmouth-Hitchcock Medical Center: Laura A. Flashman, PhD; Marc Seltzer, MD; Mary L. Hynes, RN, MPH; Robert B. Santulli, MD (past investigator); Wake Forest University Health Sciences: Kaycee M. Sink, MD, MAS; Mia Yang, MD; Akiva Mintz, MD, PhD; Rhode Island Hospital: Brian R. Ott, MD; Geoffrey Tremont, PhD; Lori A. Daiello, Pharm.D, ScM; Butler Hospital: Stephen Salloway, MD, MS; Paul Malloy, PhD; Stephen Correia, PhD; Athena Lee, PhD; UC San Francisco: Howard J. Rosen, MD; Bruce L. Miller, MD; David Perry, MD; Medical University South Carolina: Jacobo Mintzer, MD, MBA; Kenneth Spicer, MD, PhD; David Bachman, MD; St. Joseph’s Health Care: Elizabeth Finger, MD; Stephen Pasternak, MD; Irina Rachinsky, MD; John Rogers, MD; Andrew Kertesz, MD (past investigator); Dick Drost, MD (past investigator); Nathan Kline Institute: Nunzio Pomara, MD; Raymundo Hernando, MD; Antero Sarrael, MD; University of Iowa College of Medicine: Delwyn D. Miller, PharmD, MD; Karen Ekstam Smith, RN; Hristina Koleva, MD; Ki Won Nam, MD; Hyungsub Shim, MD; Susan K. Schultz, MD (past investigator); Cornell University Norman Relkin, MD, PhD; Gloria Chiang, MD; Michael Lin, MD; Lisa Ravdin, PhD; University of South Florida: USF Health Byrd Alzheimer’s Institute: Amanda Smith, MD; Christi Leach, MD; Balebail Ashok Raj, MD (past investigator); Kristin Fargher, MD (past investigator); Courtney Bodge, PhD; ADNI Part A: Leadership and Infrastructure Principal Investigator: Michael W. Weiner, MD, University of California, San Francisco; ATRI Principal Investigator and Director of Coordinating Center Clinical Core: Paul Aisen, MD, University of Southern California; Executive Committee: Michael Weiner, MD, University of California, San Francisco; Paul Aisen, MD, University of Southern California; Ronald Petersen, MD, PhD, Mayo Clinic, Rochester; Robert C. Green, MD, MPH, Brigham and Women’s Hospital/Harvard Medical School; Danielle Harvey, PhD, University of California, Davis; Clifford R. Jack, Jr, MD, Mayo Clinic, Rochester; William Jagust, MD, University of California, Berkeley; John C. Morris, MD, Washington University, St Louis; Andrew J. Saykin, PsyD, Indiana University; Leslie M. Shaw, PhD, Perelman School of Medicine, University of Pennsylvania; Arthur W. Toga, PhD, University of Southern California; John Q. Trojanowki, MD, PhD, Perelman School of Medicine, University of Pennsylvania; Psychological Evaluation/PTSD Core: Thomas Neylan, MD, University of California, San Francisco; Traumatic Brain Injury/TBI Core: Jordan Grafman, PhD, Rehabilitation Institute of Chicago, Feinberg School of Medicine, Northwestern University; Data and Publication Committee (DPC): Robert C. Green, MD, MPH, Brigham and Women’s Hospital/Harvard Medical School (chair); Resource Allocation Review Committee: Tom Montine, MD, PhD, University of Washington (chair); Clinical Core Leaders: Michael Weiner MD (core principal investigator); Ronald Petersen, MD, PhD, Mayo Clinic, Rochester (core principal investigator); Paul Aisen, MD, University of Southern California; Clinical Informatics and Operations: Gustavo Jimenez, MBS, University of Southern California; Michael Donohue, PhD, University of Southern California; Devon Gessert, BS, University of Southern California; Kelly Harless, BA, University of Southern California; Jennifer Salazar, MBS, University of Southern California; Yuliana Cabrera, BS, University of Southern California; Sarah Walter, MSc, University of Southern California; Lindsey Hergesheimen, BS, University of Southern California; San Francisco Veterans Affairs Medical Center: Thomas Neylan, MD, University of California, San Francisco; Jacqueline Hayes, University of California, San Francisco; Shannon Finley, University of California, San Francisco; Biostatistics Core Leaders and Key Personnel: Danielle Harvey, PhD, University of California, Davis (core principal investigator); Michael Donohue, PhD, University of California, San Diego; MRI Core Leaders and Key Personnel: Clifford R. Jack, Jr, MD, Mayo Clinic, Rochester (core principal investigator); Matthew Bernstein, PhD, Mayo Clinic, Rochester; Bret Borowski, RT, Mayo Clinic; Jeff Gunter, PhD, Mayo Clinic; Matt Senjem, MS, Mayo Clinic; Kejal Kantarci, Mayo Clinic; Chad Ward, Mayo Clinic; PET Core Leaders and Key Personnel: William Jagust, MD, University of California, Berkeley (core principal investigator); Robert A. Koeppe, PhD, University of Michigan; Norm Foster, MD, University of Utah; Eric M. Reiman, MD, Banner Alzheimer’s Institute; Kewei Chen, PhD, Banner Alzheimer’s Institute; Susan Landau, PhD, University of California, Berkeley; Neuropathology Core Leaders: John C. Morris, MD, Washington University, St Louis; Nigel J. Cairns, PhD, FRCPath, Washington University, St Louis; Erin Householder, MS, Washington University, St Louis; Biomarkers Core Leaders and Key Personnel: Leslie M. Shaw, PhD, Perelman School of Medicine, University of Pennsylvania; John Q. Trojanowki, MD, PhD, Perelman School of Medicine, University of Pennsylvania; Virginia Lee, PhD, MBA, Perelman School of Medicine, University of Pennsylvania; Magdalena Korecka, PhD, Perelman School of Medicine, University of Pennsylvania; Michal Figurski, PhD, Perelman School of Medicine, University of Pennsylvania; Informatics Core Leaders and Key Personnel: Arthur W. Toga, PhD, University of Southern California (core principal investigator); Karen Crawford, University of Southern California; Scott Neu, PhD, University of Southern California; Genetics Core Leaders and Key Personnel: Andrew J. Saykin, PsyD, Indiana University; Tatiana M. Foroud, PhD, Indiana University; Steven Potkin, MD, University of California, Irvine; Li Shen, PhD, Indiana University; Kelley Faber, MS, CCRC, Indiana University; Sungeun Kim, PhD, Indiana University; Kwangsik Nho, PhD, Indiana University; Initial Concept Planning & Development: Michael W. Weiner, MD, University of California, San Francisco; Karl Friedl, US Department of Defense (retired); Part B: Investigators by Site: University of Southern California: Lon S. Schneider, MD, MS; Sonia Pawluczyk, MD; Mauricio Becerra; University of California, San Diego: James Brewer, MD, PhD; Helen Vanderswag, RN; Columbia University Medical Center: Yaakov Stern, PhD; Lawrence S. Honig, MD, PhD; Karen L. Bell, MD; Rush University Medical Center: Debra Fleischman, PhD; Konstantinos Arfanakis, PhD; Raj C. Shah, MD; Wien Center: Ranjan Duara, MD (principal investigator); Daniel Varon, MD (co–principal investigator); Maria T. Greig (HP coordinator); Duke University Medical Center: P. Murali Doraiswamy, MBBS; Jeffrey R. Petrella, MD; Olga James, MD; University of Rochester Medical Center: Anton P. Porsteinsson, MD (director); Bonnie Goldstein, MS, NP (coordinator); Kimberly S. Martin, RN; University of California, Irvine: Steven G. Potkin, MD; Adrian Preda, MD; Dana Nguyen, PhD; Medical University South Carolina: Jacobo Mintzer, MD, MBA; Dino Massoglia, MD, PhD, Olga Brawman-Mintzer, MD; Premiere Research Inst (Palm Beach Neurology): Carl Sadowsky, MD; Walter Martinez, MD; Teresa Villena, MD; University of California, San Francisco: William Jagust MD; Susan Landau, PhD; Howard Rosen, MD; David Perry; Georgetown University Medical Center: Raymond Scott Turner, MD, PhD; Kelly Behan Brigid Reynolds, NP; Brigham and Women's Hospital: Reisa A. Sperling, MD; Keith A. Johnson, MD; Gad Marshall, MD; Banner Sun Health Research Institute: Marwan N. Sabbagh, MD; Sandra A. Jacobson, MD; Sherye A. Sirrel, MS, CCRC; Howard University: Thomas O. Obisesan, MD, MPH; Saba Wolday, MSc; Joanne Allard, PhD; University of Wisconsin: Sterling C. Johnson, Ph.D. J. Jay Fruehling, MA; Sandra Harding, MS; University of Washington: Elaine R. Peskind, MD; Eric C. Petrie, MD, MS; Gail Li, MD, PhD; Stanford University: Jerome A. Yesavage, MD; Joy L. Taylor, PhD; Ansgar J. Furst, PhD; Steven Chao, MD; Cornell University: Norman Relkin, MD, PhD, Gloria Chiang, MD; Lisa Ravdin, PhD; ADNI Depression Part A: Leadership and Infrastructure Principal Investigator: Scott Mackin, PhD, University of California, San Francisco; ATRI PI and Director of Coordinating Center Clinical Core: Paul Aisen, MD, University of Southern California; Rema Raman, PhD, University of Southern California; Executive Committee: Scott Mackin, PhD, University of California, San Francisco; Michael Weiner, MD, University of California, San Francisco; Paul Aisen, MD, University of Southern California; Rema Raman, PhD, University of Southern California; Clifford R. Jack, Jr, MD, Mayo Clinic, Rochester; Susan Landau, PhD, University of California, Berkeley; Andrew J. Saykin, PsyD, Indiana University; Arthur W. Toga, PhD, University of Southern California; Charles DeCarli, MD, University of California, Davis; Robert A. Koeppe, PhD, University of Michigan; Data and Publication Committee (DPC): Robert C. Green, MD, MPH, Brigham and Women’s Hospital/Harvard Medical School (chair); Erin Drake, MA, Brigham and Women’s Hospital/Harvard Medical School (director); Clinical Core Leaders: Michael Weiner, MD (core principal investigator); Paul Aisen, MD, University of Southern California; Rema Raman, PhD, University of Southern California; Mike Donohue, PhD, University of Southern California; Clinical Informatics, Operations and Regulatory Affairs: Gustavo Jimenez, MBS, University of Southern California; Devon Gessert, BS, University of Southern California; Kelly Harless, BA, University of Southern California; Jennifer Salazar, MBS, University of Southern California; Yuliana Cabrera, BS, University of Southern California; Sarah Walter, MSc, University of Southern California; Lindsey Hergesheimer, BS, University of Southern California; Elizabeth Shaffer, BS; Psychiatry Site Leaders and Key Personnel: Scott Mackin, PhD, University of California, San Francisco; Craig Nelson, MD, University of California, San Francisco; David Bickford, BA, University of California, San Francisco; Meryl Butters, PhD, University of Pittsburgh; Michelle Zmuda, MA, University of Pittsburgh; MRI Core Leaders and Key Personnel: Clifford R. Jack, Jr, MD, Mayo Clinic, Rochester (core principal investigator); Matthew Bernstein, PhD, Mayo Clinic, Rochester; Bret Borowski, RT, Mayo Clinic, Rochester; Jeff Gunter, PhD, Mayo Clinic, Rochester; Matt Senjem, MS, Mayo Clinic, Rochester; Kejal Kantarci, MD, Mayo Clinic, Rochester; Chad Ward, BA, Mayo Clinic, Rochester; Denise Reyes, BS, Mayo Clinic, Rochester; PET Core Leaders and Key Personnel: Robert A. Koeppe, PhD, University of Michigan Susan Landau, PhD, University of California, Berkeley; Informatics Core Leaders and Key Personnel: Arthur W. Toga, PhD, University of Southern California (core principal investigator); Karen Crawford, University of Southern California; S Neu, PhD, University of Southern California; Genetics Core Leaders and Key Personnel: Andrew J. Saykin, PsyD, Indiana University; Tatiana M. Foroud, PhD, Indiana University; Kelley M. Faber, MS, CCRC, Indiana University; Kwangsik Nho, PhD, Indiana University; Kelly N. Nudelman, Indiana University; Part B: Investigators by Site: University of California, San Francisco: Scott Mackin, PhD; Howard Rosen, MD; Craig Nelson, MD; David Bickford, BA; Yiu Ho Au, BA; Kelly Scherer, BS; Daniel Catalinotto, BA; Samuel Stark, BA; Elise Ong, BA; Dariella Fernandez, BA; University of Pittsburgh: Meryl Butters, PhD; Michelle Zmuda, MA; Oscar L. Lopez, MD; MaryAnn Oakley, MA; Donna M. Simpson, CRNP, MPH.
